# Combining CRISPR/Cas9 and brain imaging: from genes to molecules to networks

**DOI:** 10.1101/2021.09.10.459766

**Authors:** Sabina Marciano, Tudor Mihai Ionescu, Ran Sing Saw, Rachel Y. Cheong, Deniz Kirik, Andreas Maurer, Bernd Pichler, Kristina Herfert

## Abstract

Receptors, transporters and ion channels are important targets for therapy development in neurological diseases including Alzheimeŕs disease, Parkinsońs disease, epilepsy, schizophrenia and major depression. Several receptors and ion channels identified by next generation sequencing may be involved in disease initiation and progression but their mechanistic role in pathogenesis is often poorly understood. Gene editing and *in vivo* imaging approaches will help to identify the molecular and functional role of these targets and the consequence of their regional dysfunction on whole brain level. Here, we combine CRISPR/Cas9 gene-editing with *in vivo* positron emission tomography (PET) and functional magnetic resonance imaging (fMRI) to investigate the direct link between genes, molecules, and the brain connectome. The extensive knowledge of the *Slc18a2* gene encoding the vesicular monoamine transporter (VMAT2), involved in the storage and release of dopamine, makes it an excellent target for studying the gene networks relationships while structurally preserving neuronal integrity and function. We edited the *Slc18a2* in the substantia nigra pars compacta of adult rats and used *in vivo* molecular imaging besides behavioral, histological, and biochemical assessments to characterize the CRISPR/Cas9-mediated VMAT2 knockdown. Simultaneous PET/fMRI was performed to investigate molecular and functional brain alterations. We found that stage-specific adaptations of brain functional connectivity follow the selective impairment of presynaptic dopamine storage and release. Our study reveals that recruiting different brain networks is an early response to the dopaminergic dysfunction preceding neuronal cell loss. Our combinatorial approach is a novel tool to investigate the impact of specific genes on brain molecular and functional dynamics which will help to develop tailored therapies for normalizing brain function. The method can easily be transferred to higher-order species allowing for a direct comparison of the molecular imaging findings.

## Introduction

The brain is a network of spatially distributed but functionally and structurally interconnected regions that exhibit correlated activity over time. They communicate with each other via highly specialized neuronal connections and are organized in neuronal circuits and networks. Understanding how functional connections between regions are arranged in the healthy and diseased brain is therefore of great interest.

Resting-state functional magnetic resonance imaging (rs-fMRI) has enabled neuroscientists to delineate the level of functional communication between anatomically separated regions [1]. Rs-fMRI measures the resting-state functional connectivity (rs-FC) at high spatial and temporal resolutions based on spontaneous fluctuations of the blood oxygen level-dependent (BOLD) signal at rest, which indirectly detects neuronal activity via hemodynamic coupling [2]. Using rs-fMRI several brain resting-state networks in humans and rodents have been identified, such as the default mode and sensorimotor networks (DMN, SMN) [3–9]. Alterations of these networks are linked to neurological diseases [10, 11], and may serve as early therapeutic and diagnostic biomarkers. However, the molecular signatures related to the functional alterations in disease remain largely unknown.

Positron emission tomography (PET) provides a non-invasive tool to indirectly measure molecular changes in the brain with high specificity and sensitivity. One well-characterized example is the radioligand [^11^C]raclopride, a widely used D2/D3 receptor antagonist enabling the non-invasive determination of dopamine release and availability [12–14].

PET in combination with BOLD-fMRI has the great potential to investigate the molecular substrate of brain functional connectivity (FC), enabling the direct spatial and temporal correlation of both measurements [15–21]. In this context, we have recently shown that rs-FC is modulated by intrinsic serotonin transporter and D2/3 receptor occupancy in rats [22].

Insights into functional brain circuits and their relationships to individual phenotypes can be gained by genetic manipulations of neuronal subtypes [23]. Genome-engineering methodologies based on clustered regularly interspaced short palindromic repeats (CRISPR)/associated RNA-guided endonuclease (Cas9) represent a promising approach to unveil the influence of genes on brain circuits. CRISPR/Cas9 has enabled researchers to interrogate the mammalian DNA in a precise yet simple manner [24, 25] in several species [26–31], by editing single or multiple genomic loci *in vitro* and *in vivo* [25, 32, 33]. However, one great hurdle is the brain delivery, which must comply with effective nuclear access, while minimizing immunogenic reactions and off-target editing [34]. Despite these limitations, the potential of CRISPR/Cas9 is continuously expanding with novel nuclease variants being exploited [35–37]. Derived from *Staphylococcus aureus*, SaCas9 overcomes the packaging constraints of adeno-associated viral vectors (AAVs), allowing efficient CRISPR/Cas9 brain transfer [38–42].

Here, we use an AAV-based CRISPR/SaCas9 gene-editing approach to knock down the *Slc18a2* gene encoding the vesicular monoamine transporter 2 (VMAT2), a key protein involved in the storage and release of dopamine in the brain [43]. The extensive knowledge on *Slc18a2* makes it an excellent basis for studying the gene networks relationships. We characterize the VMAT2- mediated dopamine signaling using *in vivo* molecular imaging, behavioral, histological, and biochemical assessments. We investigate the impact of impaired VMAT2-dependent dopamine neurotransmission on the DMN and SMN using a simultaneous [^11^C]raclopride-PET/fMRI protocol. Further, we dig into the dopamine-GABA interplay using [^11^C]flumazenil PET. Our results reveal that CRISPR/SaCas9-induced synaptic dysfunction prompts early network changes, preceding motor and molecular alterations, including a regional increase in postsynaptic dopamine receptor availability. We identify a pattern of asymmetric hyperconnectivity, and internetwork synchronization, spreading from the contralateral thalamus (SMN), and prefrontal cortical regions (DMN) to the striata and hippocampi, complemented by a reduced GABA-A receptor availability.

Our findings illustrate the ability of the brain to recruit different brain networks and functionally compensate for the dopaminergic dysfunction prior to neuronal cell loss, postsynaptic changes, and motor impairment.

## Results

### *In vitro* validation of CRISPR/SaCas9-induced VMAT2 knockdown in rat primary cortical neurons

To evaluate the efficiency of the AAV-based CRISPR/SaCas9 VMAT2 knockdown in rat primary neurons, we designed AAV-SaCas9 and AAV-sgRNA targeting the first exon of the bacterial *lacZ* gene (control) or the second exon of the *Slc18a2* gene (Fig. 1a) (sgRNAs sequences are reported in Table 1). Seven days post-transduction, the protein expression level and mutation rate of the harvested genomic DNA were inspected by immunofluorescence and surveyor assay (Fig. 1b). Immunofluorescence indicated a clear reduction of VMAT2 protein expression in neurons transduced with AAV-SaCas9 and AAV-sgRNA-*Slc18a2* (Fig. 1c). We observed 20% editing for the digested DNA from neurons transduced with vectors for SaCas9 and sgRNA- *Slc18a2* (Supplementary Fig.1).

**Fig. 1.**
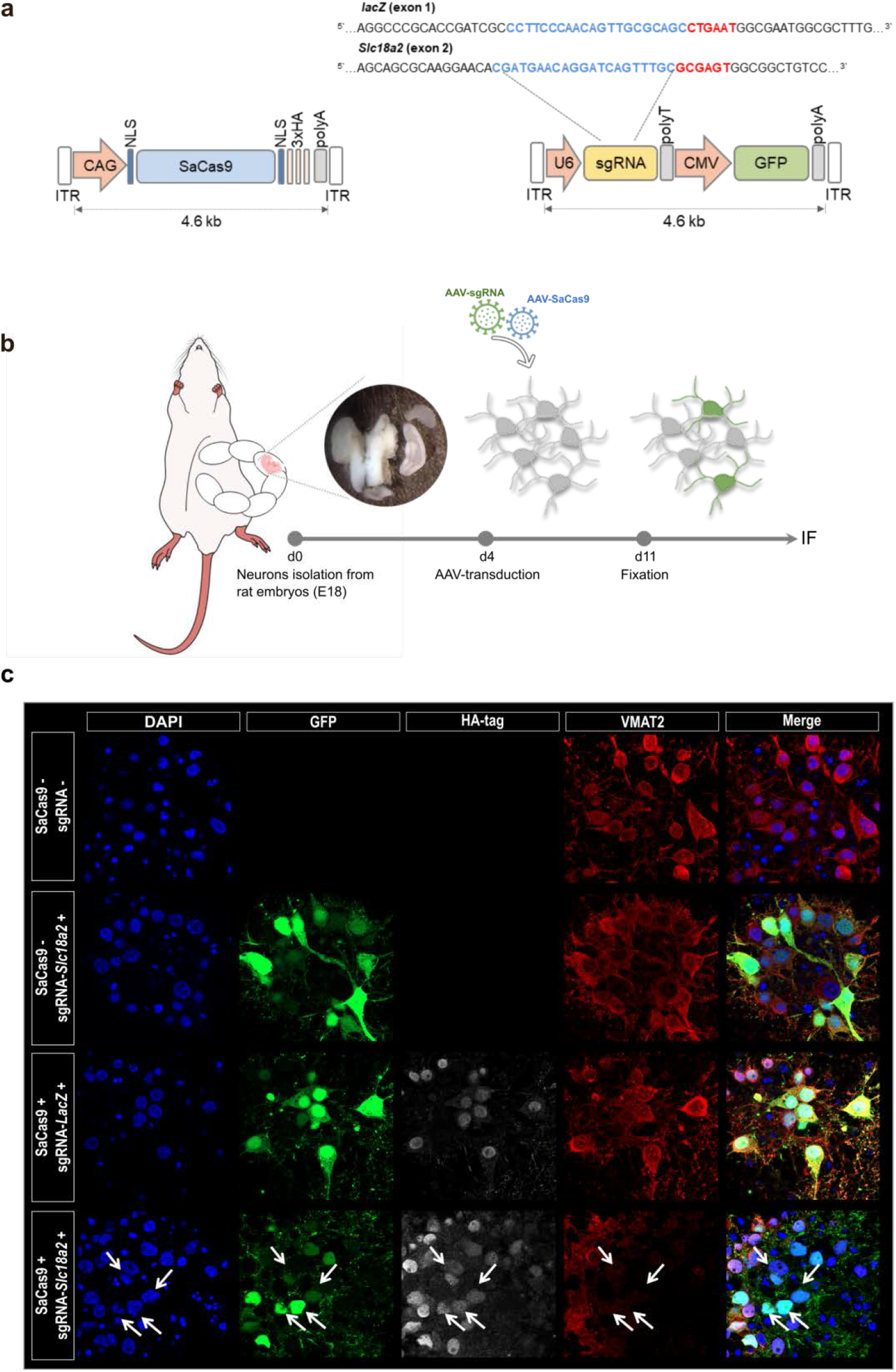
*In vitro* validation of CRISPR/SaCas9-induced VMAT2 knockdown in rat primary cortical neurons. (**a**) AAV-SaCas9 and AAV-sgRNA expression vectors. (**b**) Experimental design for primary neurons isolation and transduction. (**c**) VMAT2 immunostaining (red), nuclei labeled with DAPI (blue). GFP (green) and HA-tag (white) indicate the expression of the sgRNA and SaCas9 vectors, respectively. VMAT2 KD (arrows) is shown in neurons transduced with AAVs carrying SaCas9 and sgRNA-*Slc18a2.* KD, knockdown; ITR, inverted terminal repeat; CMV, cytomegalovirus promoter; GFP, green fluorescent protein; CAG, CMV enhancer/chicken β-actin promoter; NLS, nuclear localization signal; HA-tag, hemagglutinin tag; polyA, polyadenylation signal; polyT, polytermination signal.

**Table 1:**
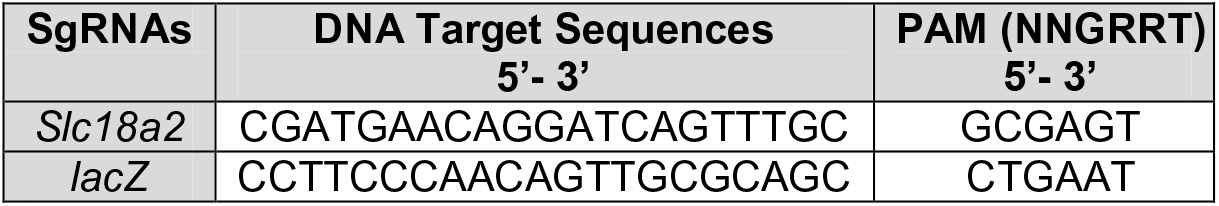
sgRNAs sequences.

### CRISPR/SaCas9-induced VMAT2 knockdown elicits postsynaptic changes but no nerve terminal loss or neuroinflammation in the adult rat brain

To test the *in vivo* efficiency of the CRISPR/SaCas9 gene-editing, we expressed SaCas9 and sgRNA targeting *Slc18a2* to knock down the VMAT2, or targeting *lacZ* as control, by AAV-mediated gene transfer into the right substantia nigra pars compacta (SNc). DPBS was injected into the left SNc. [^11^C]Dihydrotetrabenazine (DTBZ) PET imaging was performed 8 – 10 weeks post-injection to quantify VMAT2 expression in the striatum (Fig. 2a). [^11^C]DTBZ binding was decreased by 30% (0-62%) in the right striatum of rats where the VMAT2 was knocked down in comparison to the contralateral striatum. No changes of [^11^C]DTBZ binding were observed in the contralateral striatum, as [^11^C]DTBZ binding did not differ between the left striatum of rats injected with sgRNA targeting *lacZ* and rats injected with sgRNA targeting *Slc18a2* (Fig. 2b,c).

**Fig. 2.**
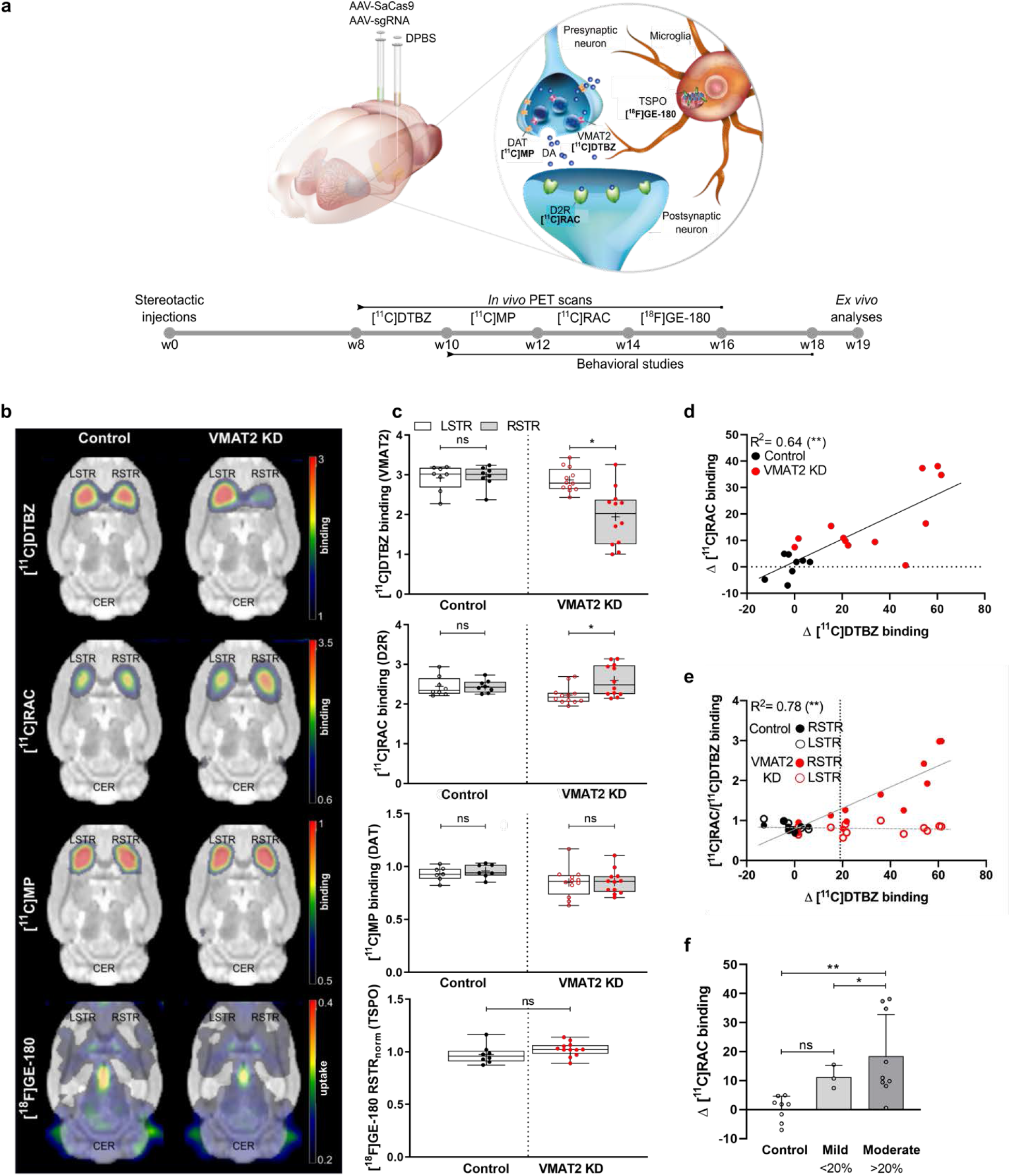
CRISPR/SaCas9-induced VMAT2 knockdown elicits postsynaptic changes but no nerve terminal loss or neuroinflammation in the adult rat brain. (**a**) Schematic illustration of the experimental design. (**b**) Mean binding potential and uptake maps of control and VMAT2 KD rats co-registered to a rat brain atlas. (**c**) Binding potential values of individual control and VMAT2 KD rats in the left and right striatum. For [^18^F]GE-180 uptake values normalized to the left striatum are shown. (**d**) A strong correlation between Δ [^11^C]RAC and Δ [^11^C]DTBZ binding is shown. (**e**) Ratio of striatal [^11^C]RAC and [^11^C]DTBZ binding shows prominent [^11^C]RAC changes when a threshold of ∼ 20% Δ [^11^C]DTBZ binding is reached. This threshold was set to separate the VMAT2 KD rats into *mild* and *moderate*. (**f**) *Mild* and *moderate* rats could be differentiated based on the postsynaptic changes. *\*P*< 0.01, \**\*P*< 0.001, Bonferroni-Sidak corrected. Data are shown as boxplot with the median value (central mark), the mean value (plus sign), interquartile range (boxes edges), and the extreme points of the distribution (whiskers). Control rats n= 8; VMAT2 KD rats n= 12. *Mild*: Δ [^11^C]DTBZ binding < 20%; *Moderate*: Δ [^11^C]DTBZ binding ≥ 20%. [^11^C]MP, [^11^C]methylphenidate; [^11^C]RAC, [^11^C]raclopride; LSTR, left striatum; RSTR, right striatum; CER, cerebellum; KD, knockdown; DAT, dopamine transporter; TSPO, translocator protein.

We further evaluated changes of dopamine availability in the striatum using [^11^C]raclopride, which competes with dopamine for the same binding site at the D2 receptor (D2R) [13]. After 12 – 14 weeks following CRISPR/SaCas9-induced VMAT2 knockdown in nigrostriatal neurons, we observed 17% increased binding of [^11^C]raclopride in the right striatum of VMAT2 knockdown rats and no changes in control rats (Fig. 2b,c), indicating a reduction of synaptic dopamine levels and/or compensatory changes of D2R expression at postsynaptic medium spiny neurons. A larger VMAT2 knockdown led to lower dopamine levels in the striatum and thus to higher D2R binding (Fig. 2d). To explore the threshold at which the observed postsynaptic changes occur, we calculated the [^11^C]raclopride/[^11^C]DTBZ binding ratio for the right and left striatum. The ratio remained close to 1 in the DPBS-injected striatum and control rats, indicating no substantial difference between the two hemispheres. In contrast, VMAT2 knockdown rats displayed large [^11^C]raclopride binding changes when the level of VMAT2 knockdown was ∼ 20%. From this point, a prominent increase in D2R binding was observed in the right striatum (Fig. 2e). Therefore, this threshold was set to split the rats into *mild* (< 20%) and *moderate* (≥ 20%). Notably, [^11^C]raclopride PET imaging was able to discriminate between different degrees of synaptic dysfunction, classified from [^11^C]DTBZ binding changes (Fig. 2f).

We inspected the integrity of dopaminergic nerve terminals and the occurrence of neuroinflammation in the striatum after the CRISPR/SaCas9-induced VMAT2 knockdown. [^11^C]methylphenidate PET imaging of the dopamine transporter and [^18^F]GE-180 PET imaging of the translocator protein, which is overexpressed on activated microglia, was performed. CRISPR/SaCas9-induced VMAT2 knockdown did neither alter [^11^C]methylphenidate binding, nor [^18^F]GE-180 uptake (Fig. 2b,c).

### CRISPR/SaCas9-induced VMAT2 knockdown impairs motor function

To explore the motor consequences of the CRISPR/SaCas9-induced VMAT2 knockdown, we performed several behavioral tests (Fig. 3a).

**Fig. 3.**
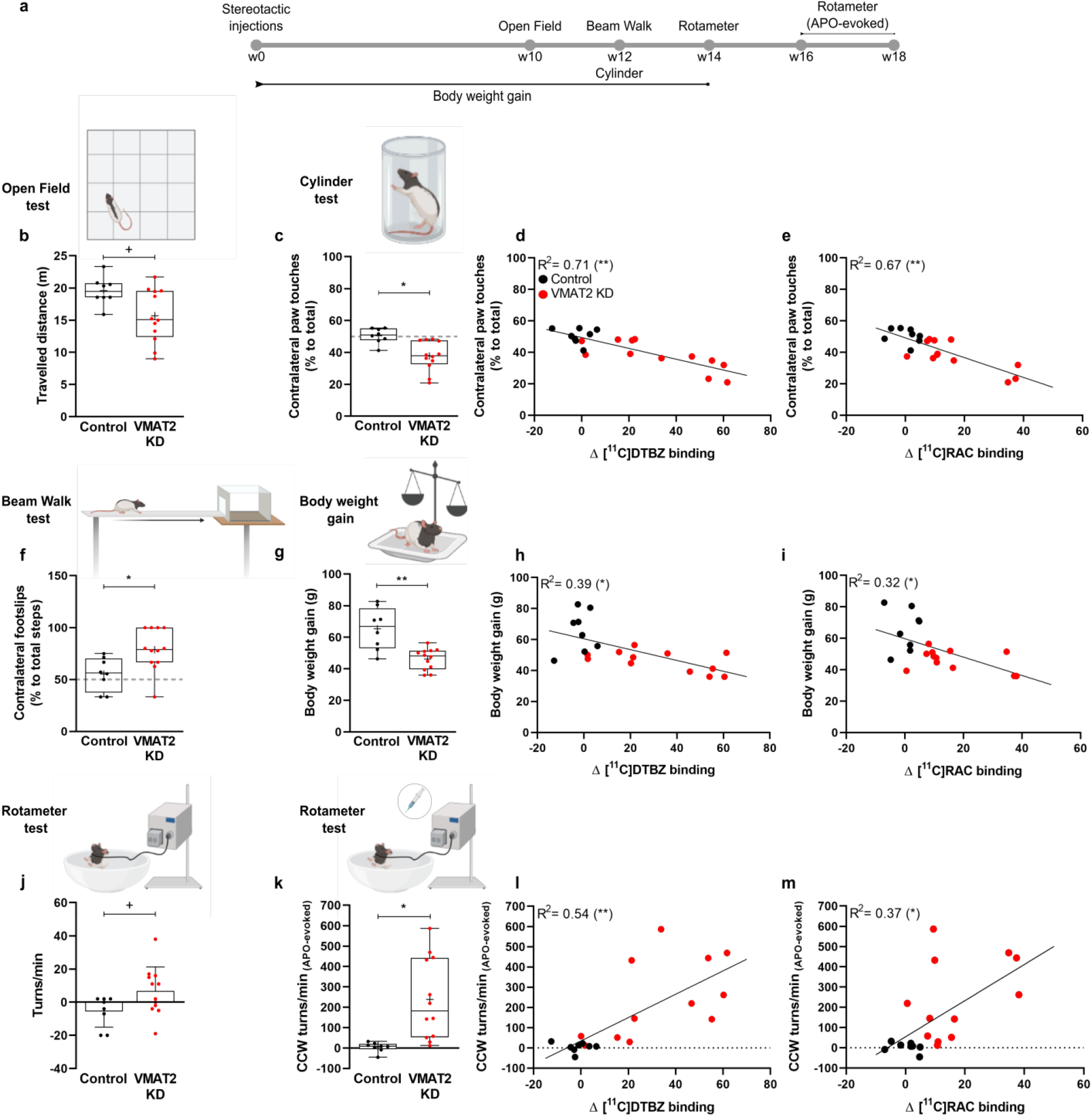
CRISPR/SaCas9-induced VMAT2 knockdown impairs motor function. (**a**) Schematic illustration of the behavioral tests. (**b**) In the open field test, the distance travelled (m) by VMAT2 KD rats was reduced. (**c**) Cylinder test. VMAT2 KD rats showed a reduction in the contralateral paw touches, compared with controls. Rats performance in the cylinder test strongly correlated with VMAT2 expression changes (Δ [^11^C]DTBZ binding) and corresponding changes in dopamine availability (Δ [^11^C]RAC binding) (**d,e**). (**f**) In the beam walk test, VMAT2 KD rats displayed a higher number of footslips to the left contralateral side compared with control rats. (**g**) Body weight assessment 14 weeks after CRISPR/SaCas9 gene-editing showed reduced body weight gain in VMAT2 KD compared with control rats. (**h,i**) Body weight gain correlated with VMAT2 expression changes (Δ [^11^C]DTBZ binding) and corresponding changes in dopamine availability (Δ [^11^C]RAC binding). (**j**) Spontaneous rotation in a novel spherical environment showed increased CW rotations in VMAT2 KD rats compared with control rats. (**k**) Apomorphine-evoked rotational behavior. VMAT2 KD rats displayed a higher number of CCW rotations compared with control rats in the rotameter test. (**l,m**) Apomorphine-evoked rotations exhibited a strong correlation with VMAT2 expression changes (Δ [^11^C]DTBZ binding) and changes in dopamine availability (Δ [^11^C]RAC binding). Data are shown as boxplot with the median value (central mark), the mean value (plus sign), interquartile range (boxes edges), and the extreme points of the distribution (whiskers). ^+^*P*< 0.05, *\*P*< 0.01, *\*\*P*< 0.001. Control rats n= 8; VMAT2 KD rats n= 12. CCW, counter-clockwise; APO, apomorphine; KD, knockdown; [^11^C]RAC, [^11^C]raclopride. Illustrations in the figure were created with BioRender.com.

We observed a reduction in the locomotor activity of VMAT2 knockdown rats in the open field test (Fig. 3a), but no correlation to VMAT2 expression changes (Δ [^11^C]DTBZ binding), or dopamine availability (Δ [^11^C]raclopride binding) (Supplementary Fig. 2a,b).

Next, we evaluated the forelimb akinesia using the cylinder test. VMAT2 knockdown rats displayed a preference for the right forepaw, while control rats equivalently used their right and left forepaw (Fig. 3c). Paw use alterations correlated highly with VMAT2 knockdown (Δ [^11^C]DTBZ binding), and dopamine availability (Δ [^11^C]raclopride binding) (Fig. 3d,e).

To further examine differences in motor function, coordination, and balance, rats underwent the beam walk test. VMAT2 knockdown rats stumbled with higher frequency to the left side, while control rats displayed equal chances to slip in each direction (Fig. 3f). However, no correlations between gait alterations and VMAT2 knockdown (Δ [^11^C]DTBZ binding), or dopamine availability (Δ [^11^C]raclopride binding) were found (Supplementary Fig. 2c,d).

As previous studies suggest that body weight changes reflect striatal dopamine depletion [44], we inspected the impact of the VMAT2 knockdown on the rats’ body weight gain. VMAT2 knockdown rats exhibited a 30% reduction in their gained weight over a period of 14 weeks, compared with controls (Fig. 3g). Body weight gain correlated with changes in VMAT2 expression (Δ [^11^C]DTBZ binding), and dopamine availability (Δ [^11^C]raclopride binding) (Fig. 3h,i).

To assess the rotational behavior, we performed the rotameter test with and without apomorphine administration. In the spontaneous rotation test, VMAT2 knockdown rats displayed a higher number of ipsilateral net turns compared with control rats (Fig. 3j). The number of turns did not correlate with VMAT2 expression changes (Δ [^11^C]DTBZ binding), and changes in dopamine availability (Δ [^11^C]raclopride binding) (Supplementary Fig. 2e,f). Apomorphine-induced rotations to the contralateral side were higher in VMAT2 knockdown rats compared with control rats (Fig. 3k), and correlated with changes in VMAT2 expression (Δ [^11^C]DTBZ binding) and dopamine availability (Δ [^11^C]raclopride binding) (Fig. 3l,m).

### *Ex vivo* validation of the CRISPR/SaCas9-induced VMAT2 knockdown

Using immunofluorescence, we confirmed the concomitant expression of SaCas9 and *Slc18a2*– targeting sgRNA 19 weeks post-transduction, and a corresponding decrease of VMAT2 expression in the SNc of the VMAT2 knockdown group (Fig. 4b).

**Fig. 4.**
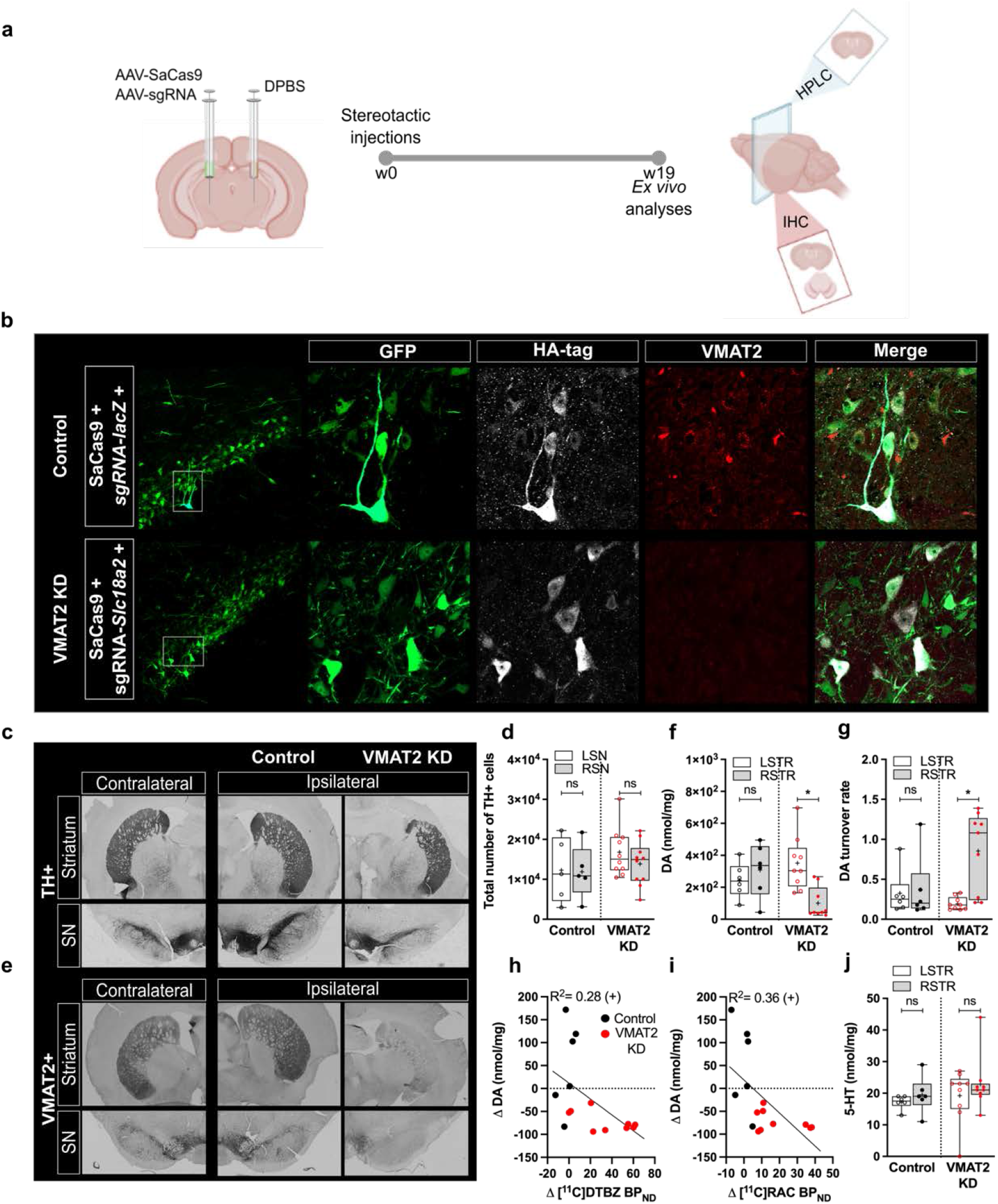
*Ex vivo* validation of the CRISPR/SaCas9-induced VMAT2 knockdown. (**a**) Schematic illustration of the *ex vivo* analyses. (**b**) Immunofluorescence of nigral sections of control and VMAT2 KD rats confirmed the concomitant expression of SaCas9 and sgRNA. Staining for GFP (AAV-sgRNAs, green), HA- tag (AAV-SaCas9, white), and VMAT2 expression (red) for two exemplary rats is shown. VMAT2 expression was largely reduced in the SNc of VMAT2 KD rats. (**c,d**) TH expression in the ipsilateral and contralateral striatum and SNc of VMAT2 KD (n= 10) and control (n= 5) rats evidenced no cell loss. (**e**) VMAT2 immunohistochemistry in the SNc and striatum confirmed large protein reduction in the ipsilateral hemisphere of VMAT2 KD rats. (**f**) Striatal dopamine, normalized to total protein concentration, was reduced in the ipsilateral striatum of VMAT2 KD (n= 9), but not control (n= 6) rats. This reduction was paralleled by increased metabolic outcome (**g**). (**h,i**) Dopamine changes correlated with the VMAT2 KD extent and postsynaptic changes, deducted from [^11^C]DTBZ and [^11^C]RAC, respectively. (**j**) 5-HT content was unaltered in the striata of control (n= 6) and VMAT2 KD (n= 9) rats. Data are shown as boxplot with the median value (central mark), the mean value (plus sign), interquartile range (boxes edges), and the extreme points of the distribution (whiskers). ^+^*P*< 0.05, *\*P*< 0.01. GFP, green fluorescent protein; HA-tag, hemagglutinin tag; KD, knockdown; TH, tyrosine hydroxylase; SN, substantia nigra; STR, striatum; [^11^C]RAC, [^11^C]raclopride; DA, dopamine; 5-HT, serotonin. Metabolites’ and neurotransmitters’ striatal content is reported in Table 3. Illustrations in the figure were created with BioRender.com.

Immunohistochemistry revealed no changes in tyrosine hydroxylase (TH) expression levels in striatum and SN in both groups (Fig. 4c,d), and confirmed the reduction of VMAT2 expression in the right striatum and SN in the knockdown group (Fig. 4e).

Biochemical analysis showed a large reduction of dopamine, paralleled by an increased ratio of metabolites (DOPAC, HVA) to dopamine, in the right striatum of VMAT2 knockdown rats (Fig. 4f,g). The reduced dopamine content correlated with the *in vivo* VMAT2 expression (Δ [^11^C]DTBZ BP_ND_) and postsynaptic changes (Δ [^11^C]RAC BP_ND_) (Fig. 4h,i). Additionally, serotonin was unchanged in the striata of VMAT2 knockdown and control rats, suggesting dopamine nigrostriatal pathway specificity (Fig. 4j) (Metabolites’ and neurotransmitters’ striatal levels are reported in Table 3).

**Table 2:** Brain regions included in the Paxinos rat brain atlas, including their respective volumes and abbreviations.

| Brain region (ROI) | Hemisphere | ROI volume [mm<sup>3</sup>] | Abbreviation |
| --- | --- | --- | --- |
| Striatum | left | 43.552 | STR |
|  | right |  |  |
| Cingulate Cortex | left | 14.480 | CgC |
|  | right |  |  |
| Medial Prefrontal Cortex | left | 6.304 | mPFC |
|  | right |  |  |
| Motor Cortex | left | 32.608 | MC |
|  | right |  |  |
| Orbitofrontal Cortex | left | 18.936 | OFC |
|  | right |  |  |
| Parietal Cortex | left | 7.632 | PaC |
|  | right |  |  |
| Retrosplenial Cortex | left | 18.920 | RSC |
|  | right |  |  |
| Somatosensory Cortex | left | 71.600 | SC |
|  | right |  |  |
| Hippocampus | left | 25.064 | Hipp |
|  | right |  |  |
| Thalamus | left | 30.712 | Th |
|  | right |  |  |

**Table 3:** Dopamine, 3,4-Dihydroxyphenylacetic acid (DOPAC), homovanillic acid (HVA), and serotonin (5-HT) striatal content (nmol/mg) determined by HPLC.

| Neurotransmitter<br>or<br>Metabolite | nmol/mg<br>(mean ± SD) |  |
| --- | --- | --- |
| DA | Control | VMAT2 KD |
| STR-L | 244 ± 109 | 351 ± 168 |
| STR-R | 308 ± 166 | 100 ± 97 |
| DOPAC | Control | VMAT2 KD |
| STR-L | 34 ± 7 | 34 ± 11 |
| STR-R | 38 ± 4 | 27 ± 5 |
| HVA | Control | VMAT2 KD |
| STR-L | 24 ± 5 | 25 ± 5 |
| STR-R | 26 ± 6 | 19 ± 4 |
| 5-HT | Control | VMAT2 KD |
| STR-L | 17 ± 2 | 19 ± 8 |
| STR-R | 19 ± 6 | 23 ± 9 |

### Increased resting-state functional connectivity after CRISPR/SaCas9-induced VMAT2 knockdown

As multiple lines of evidence suggest a broader role of dopamine in the dynamic reconfiguration of brain networks [17, 45, 46], we next investigated the impact of unilateral dopamine depletion on brain rs-FC. A second cohort of rats underwent longitudinal simultaneous [^11^C]raclopride-PET/BOLD-fMRI scans at baseline and 8 - 14 weeks after CRISPR/SaCas9-induced VMAT2 knockdown (Fig. 5a). [^11^C]DTBZ PET scans and behavioral analysis confirmed previous findings in the first cohort, that is, an efficient depletion of the VMAT2 (20% decrease of [^11^C]DTBZ binding) (Supplementary Fig. 3a), paralleled by motor disturbances in the cylinder test (Supplementary Fig. 3e-g). In line with the findings of cohort 1, dopamine availability was decreased (10% increase in [^11^C]raclopride binding) and correlated to the extent of the VMAT2 knockdown (Supplementary Fig. 3b,c). An increase in D2R binding was observed in the right striatum when the level of VMAT2 knockdown reached ∼ 20% (Supplementary Fig. 3d), enabling a subdivision into *mild* (< 20%) and *moderate* rats (≥ 20%).

**Fig. 5.**
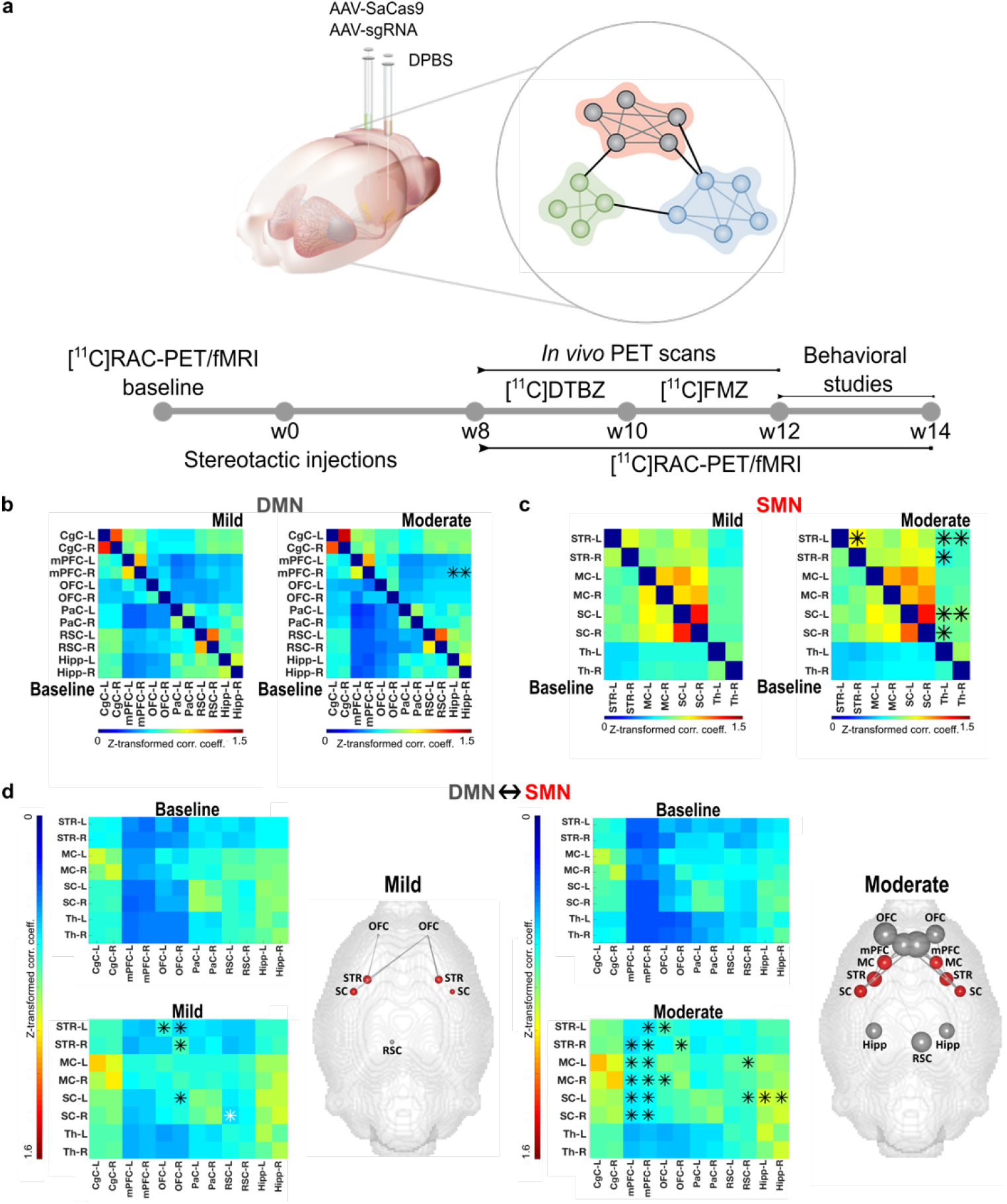
Increased resting-state functional connectivity after CRISPR/SaCas9-induced VMAT2 knockdown. (**a**) Schematic illustration of the experimental design. Group level correlation matrices of the DMN (**b**) and SMN (**c**) at baseline and after CRISPR/SaCas9-targeting for rats with a *mild* (left panel) and *moderate* (right panel) VMAT2 KD. In the *moderate* KD group rs-FC was increased between the right mPFC and right and left Hipp in the DMN (**b**) and between the contralateral SC and right and left Th, as well as between the left STR and right and left Th, in the SMN (**c**). (**d**) Internetwork rs-FC changes in the *mild* KD group indicated increased rs-FC between anterior regions of the DMN and the SMN (between the left OFC and STR, and between the right OFC and the left SC, and the right and left STR). Conversely, rs- FC was decreased between regions of the posterior DMN and the SMN (between the left RSC and right SC). In the *moderate* KD group, DMN-SMN rs-FC was increased. Brain graphs, right to the matrices, illustrate the nodes and edges (raw values) that demonstrated rs-FC changes to baseline (%). *\*P*< 0.01. *Mild*: Δ [^11^C]DTBZ binding < 20%, n= 13; *Moderate*: Δ [^11^C]DTBZ binding ≥ 20%, n= 10. KD, knockdown; DMN, default-mode network; SMN, sensorimotor network. mPFC, medial prefrontal cortex; Hipp, hippocampus; SC, somatosensory cortex; Th, thalamus; STR, striatum; OFC, orbitofrontal cortex; RSC, retrosplenial cortex. Abbreviations of brain regions considered for the analysis of the fMRI data, including their respective volumes, are reported in Table 2.

We next assessed the occurrence of rs-FC changes in DMN and SMN. Our analysis focused on identifying early biomarkers of *mild* dysfunction and patterns of spreading of synaptic dysfunction. Figure 5b,c illustrates intraregional rs-FC group-level correlation matrices at baseline and after VMAT2 knockdown in *mild* and *moderate* rats for the DMN and SMN, respectively. We observed within-network rs-FC changes in rats with *moderate* VMAT2 knockdown, in both DMN and SMN. Rats of the *mild* knockdown group revealed rs-FC changes up to 20%, in prefrontal cortical regions of the DMN, and between the left thalamus (Th) and somatosensory cortex (SC) in the SMN. However, these data need to be carefully interpreted as they did not survive a more stringent *P* value selection (\**P*< 0.01) (Supplementary Fig.4a,b) (*P* values are reported in Supplementary Tables 1,3).

Rats with *moderate* VMAT2 knockdown exhibited a 60% increase in rs-FC within the right medial prefrontal cortex (mPFC) and the right and left hippocampus (Hipp) (Fig. 5b) (*P* values are reported in Supplementary Table 2).

FC increase between the left thalamus and somatosensory cortex in the SMN doubled to 34% in rats with *moderate* VMAT2 knockdown and extended throughout the left and right thalamus and striatum (STR), respectively (Fig. 5c) (*P* values are reported in Supplementary Table 4).

Moreover, we inspected rs-FC changes between the DMN and SMN at baseline and after the CRISPR/SaCas9-induced VMAT2 knockdown.

Figure 5d illustrates internetwork rs-FC correlation matrices in rats with *mild* (left panel) and *moderate* (right panel) VMAT2 knockdown. Brain graphs display the nodes and edges (raw values) that demonstrated internetwork rs-FC changes to baseline (%). Strikingly, large alterations between DMN and SMN were already observable in the *mild* VMAT2 knockdown group. Rats presented opposite rs-FC changes between regions of the anterior/posterior DMN and the SMN, compared with baseline. A 30 to 60% increase in rs-FC was observed between regions of the anterior DMN and the SMN. Specifically, rs-FC increased between the right orbitofrontal cortex (OFC) and striatum bilaterally and the contralateral somatosensory cortex, and between the contralateral orbitofrontal cortex and striatum. Instead, a 20% decrease in rs-FC was found between regions of the posterior DMN and the SMN. Specifically, rs-FC decreased between the left retrosplenial cortex (RSC) and right somatosensory cortex (Fig. 5d, left panel) (*P* values are reported in Supplementary Table 5).

Rats with a *moderate* VMAT2 knockdown presented increased rs-FC between regions of the anterior/posterior DMN and the SMN, compared with baseline. Of particular note, internetwork rs- FC changes were not found between the regions of the posterior DMN and the SMN that showed decreased rs-FC in rats with *mild* VMAT2 knockdown. Moreover, between-network rs-FC increase extended to other regions. A 60 to 80% increase in rs-FC was found between the medial prefrontal cortex and the right striatum, and the motor (MC) and somatosensory cortex bilaterally. FC increased by more than 20% between the hippocampi and contralateral somatosensory cortex. (Fig. 5d, right panel) (*P* values are reported in Supplementary Table 6). Notably, between-network rs-FC changes did not involve the thalamus, which connectivity was however altered within the SMN.

Further, we examined how the rs-FC changes to baseline correlated between the *mild* and *moderate* groups (Supplementary Fig. 5a). Group level intraregional and internetwork rs-FC changes to baseline (%) correlated linearly between the two groups (Supplementary Fig. 5b-d). Node correlation analysis indicated a linear increase in the magnitude of the rs-FC changes to baseline in the hippocampi (Supplementary Fig. 5e), cingulate cortices (Supplementary Fig. 5f), and contralateral, but not ipsilateral, thalamus (Supplementary Fig. 5g). Our data suggest a similarity in the pattern of the intraregional and internetwork rs-FC changes between *mild* and *moderate* VMAT2 knockdown rats and a linear relationship between the magnitude of the rs-FC changes.

To complement the results of the intraregional and internetwork rs-FC, we evaluated changes in regional mean connection distances. Network-wise graph theoretical analysis on node level was paralleled by whole-brain connection-wise analysis to identify the nodes that were significantly altered in rats with *mild* and *moderate* VMAT2 knockdown, compared with baseline, for the DMN (Supplementary Fig. 6a,b) and SMN (Supplementary Fig. 6c,d). Briefly, the network organization did not change in rats with *mild* VMAT2 knockdown, compared with baseline, as changes in the global mean connection distance were not found in DMN (Supplementary Fig. 6a) nor SMN (Supplementary Fig. 6c). Interestingly, in rats with *moderate* VMAT2 knockdown network organization changes did not influence regions of the DMN (Supplementary Fig. 6b), but occurred in the contralateral striatum and thalamus (Supplementary Fig. 6d) (*P* values are reported in Supplementary Table 7).

Collectively, rs-FC results highlight lateralized effects in the SMN, as opposed to the symmetric recruitment of DMN regions.

### CRISPR/SaCas9-induced VMAT2 knockdown alters GABA signaling

Besides dopamine, dopaminergic neurons co-release GABA via the VMAT2 [47, 48]. To investigate if GABA neurotransmission is altered following the VMAT2 knockdown, we performed additional [^11^C]flumazenil PET scans 10 – 12 weeks after the CRISPR/Cas9-editing, and quantified the GABA-A binding in regions of the DMN and SMN (Fig. 6). In *mild* VMAT2 knockdown rats, we observed a decrease of [^11^C]flumazenil binding in the ipsilateral parietal cortex (PaC) (14%), hippocampus (4%), and somatosensory cortex (9%) (Fig. 6a,b). In *moderate* VMAT2 knockdown rats we observed a decrease of [^11^C]flumazenil binding in the ipsilateral parietal cortex (11%), and somatosensory cortex (6%) (Fig. 6a,c). Our data indicate that [^11^C]flumazenil binding was altered regardless of the VMAT2 knockdown extent. This was further evidenced by the lack of correlation between [^11^C]DTBZ and [^11^C]flumazenil binding changes in the target regions (linear regression data not shown).

**Fig. 6.**
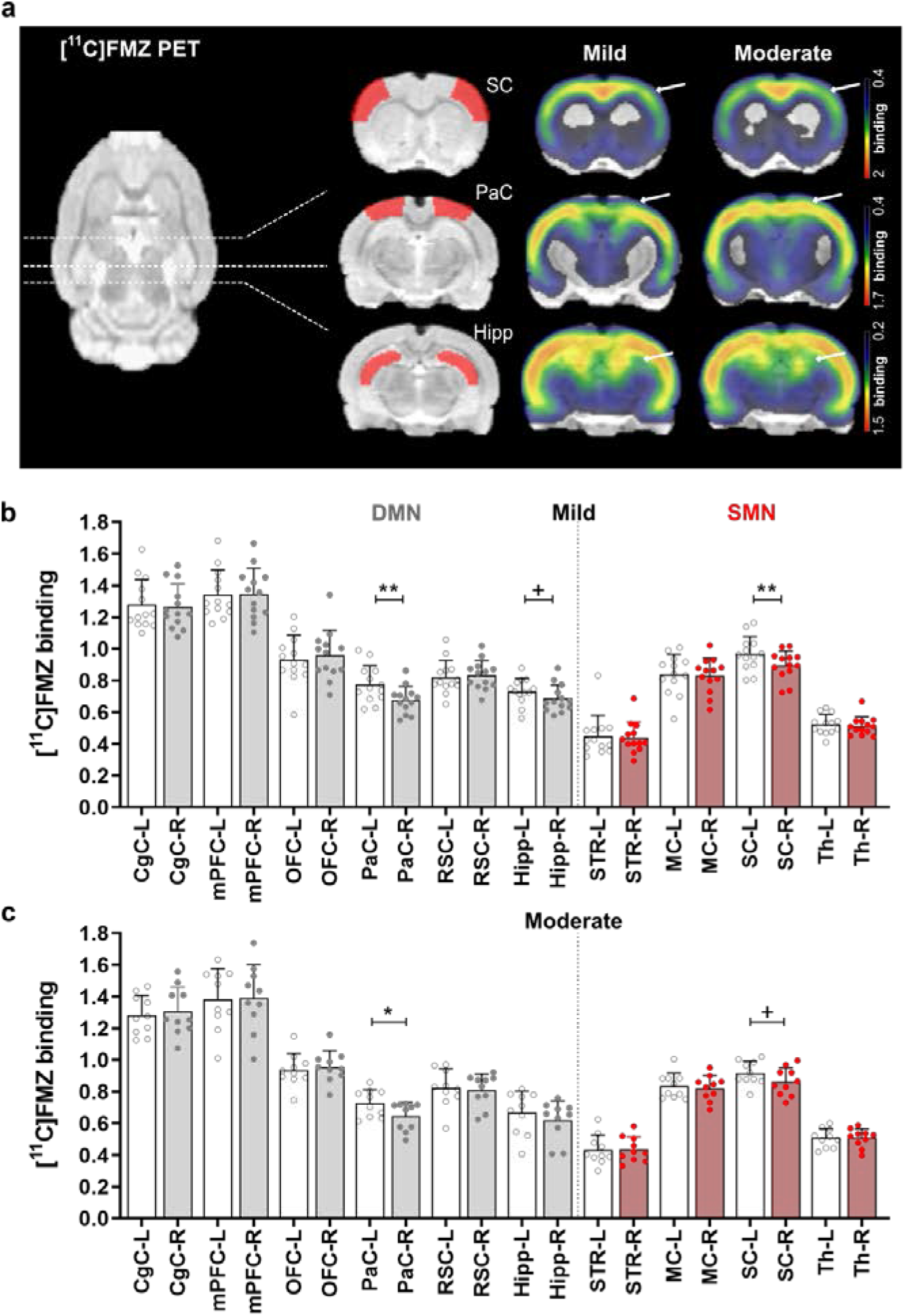
CRISPR/SaCas9-induced VMAT2 knockdown alters GABA signaling. (**a**) [^11^C]FMZ mean binding potential maps of *mild* and *moderate* VMAT2 KD rats co-registered to a rat brain atlas. Arrows and ROIs in coronal sections indicate brain regions of the DMN and SMN with altered [^11^C]FMZ binding. [^11^C]FMZ binding potentials from VOI-based analysis in DMN and SMN regions of rats with *mild* (**b**) and *moderate* (**c**) VMAT2 KD. [^11^C]FMZ binding was decreased in the right PaC, SC, and Hipp of *mild* KD rats (**b**) and right PaC and SC of *moderate* KD rats (**c**). *^+^P*< 0.05, *\*P*< 0.01, *\*\*P*< 0.001, Bonferroni-Sidak corrected. Data are shown as mean ± SD. *Mild*: Δ [^11^C]DTBZ binding < 20%, n= 13; *Moderate*: Δ [^11^C]DTBZ binding ≥ 20%, n= 10. KD, knockdown; [^11^C]FMZ, [^11^C]flumazenil; DMN, default-mode network; SMN, sensorimotor network. PaC, parietal cortex; Hipp, hippocampus; SC; somatosensory cortex. Abbreviations of brain regions, including their respective volumes, are reported in Table 2.

## Discussion

Here, we show the potential of combining CRISPR/Cas9 gene-editing with molecular and functional brain imaging to identify early adaptations of brain circuits in response to targeted gene and protein modulations. Using CRISPR/SaCas9, we knocked down the *Slc18a2* gene, encoding the VMAT2, which plays a key role in the storage and release of dopamine in response to neuronal activity [43]. The CRISPR-mediated knockdown allowed us to investigate the VMAT2- dependent dopamine signaling in the striatum, while structurally preserving neuronal integrity. [^18^F]GE-180 results suggest that glial activation is not the source of dopaminergic synaptic dysfunction, and exclude the occurrence of inflammatory responses arising from the surgical procedure and the chosen AAV-serotype, which could influence our readout, in line with recent reports [49]. Our data reveal that the targeted gene knockdown in the SNc leads to an expected reduction of dopamine release in the striatum, paralleled by [^11^C]raclopride binding changes. It is conceivable that the observed postsynaptic changes are the result of an increase in binding due to the reduced dopamine concentration in the striatum. Yet, several studies have revised this notion [50, 51]. In this regard, our results of the drug-evoked rotational behavior are in better agreement with a D2R compensatory upregulation. Indeed, supersensitivity to apomorphine in rats with nigrostriatal lesions or VMAT2 knockout is accompanied by an increase in striatal D2R binding sites, but no change in affinity [52–57]. Further, the observed striatal increase in [^11^C]raclopride binding is independent of presynaptic nerve terminal loss and occurs in response to a ∼ 20% decrease of [^11^C]DTBZ binding. This confirms, in line with our [^11^C]methylphenidate results, that [^11^C]raclopride can be used to delineate postsynaptic changes in the absence of dopamine transporter-mediated compensation, triggered by neuronal loss. Accordingly, increased [^11^C]raclopride binding is observed in the early but not later stages of Parkinson’s disease [58], characterized by severe neuronal cell loss, and dopamine transporter changes (> 50%) [59].

Consistently, rats with severe denervation (> 75%) present earlier mitigation by the dopamine transporter, followed by D2R binding changes [60]. Hence, with our method, it is feasible to study the consequences of synaptic dopamine dysfunction without compensations triggered by neuronal cell loss. Moreover, the observed [^11^C]raclopride and [^11^C]DTBZ correlations to the motor behavior highlight that [^11^C]raclopride binding remains at control levels as long as synaptic dopamine levels are sufficient to maintain adequate motor function. Motor disturbances strongly correlate to pre and postsynaptic changes if movements of the forelimbs, but not whole-body, are considered, in agreement with earlier observations in dopamine-depleted rats [44]. Depletion of VMAT2 resulted in reduced dopamine tissue levels in the ipsilateral striatum, nicely merging with the *in vivo* data. Moreover, metabolite analysis suggested that due to the lack of VMAT2- mediated storage in presynaptic vesicles, dopamine is quickly converted. The increased metabolism might as well be a possible compensatory mechanism consequent to the VMAT2 knockdown, reflecting actions that residual nigrostriatal neurons undertake to maintain dopamine homeostasis, as already speculated by others [53, 61].

To elucidate the role of VMAT2 in locomotion, reward, and Parkinson’s disease, several investigators have deleted its coding gene in mice [62–65]. Besides the costly and time-consuming breeding, the gene knockout was not selective for dopaminergic neurons, resulting in the appearance of anxiety and depressive behavior phenotypes [66]. CRISPR/Cas9-editing overcomes these limitations, allowing gene-editing in adult and aged animals and avoiding compensatory changes occurring at early developmental stages. Since its discovery, only two studies have successfully applied CRISPR/Cas9 in the rat brain [40, 67], where gene-editing has been difficult to adapt. Rats are particularly advantageous for imaging studies due to the larger brain size and limited spatial resolution and sensitivity of preclinical scanners [68]. Here, by inducing a *mild* to *moderate* gene knockdown, we could investigate early to late resting-state brain network adaptations prompted by presynaptic dysfunction. We show that the selective impairment of presynaptic dopamine storage and release is followed by rs-FC alterations within and between the DMN and SMN. Our results confirm previous findings that the DMN, associated with ideation and mind wandering [69], and the SMN, involved in sensory processing and motor function [70], do not function in isolation from each other, but rather synchronize [71]. The observed internetwork synchronization may reflect compensatory brain reorganization, as already speculated by others [72, 73]. We also identified enhanced intranetwork rs-FC in the DMN and SMN. Rs-FC changes were observed in prefrontal cortical regions, hippocampus, thalamus, and striatum. Our data parallel previous findings of cortico-striato-thalamic hyperconnectivity in decreased dopamine transmission states [74–77]. In line, increased synchronous neural oscillatory activity and functional coupling in the basal ganglia and its associated networks have been observed in Parkinson’s disease [78–83]. The increase of cortico-striatal FC could in part be due to dysfunctions of multiple tonic inhibitory gate actions of D2R [84]. Increased FC across the thalamus and prefrontal cortex has been reported in drug-treated Parkinson’s disease patients [85, 86], potentially indicating functional compensation, as the brain recruits additional anatomical areas to aid in restoring cognitive processes. This might as well explain the engagement of the hippocampus, functionally connected with DMN cortical regions [87, 88]. In this regard, research has shown that the hyperconnectivity of brain circuits is a common response to neurological dysfunction, and may reflect a protective mechanism to maintain normal brain functioning [89]. Such a mechanism has been proposed in Parkinson’s disease, mild cognitive impairment, and Alzheimer’s disease [90–93]. Collectively, our findings support this model and indicate a reorganization of brain networks that adapt to the synaptic dysfunction through enhanced interregional synchrony. Recruiting alternative brain regions may be an early response to the dysfunction preceding neuronal cell loss and motor impairment. Interestingly, brain connectome adaptations occurred symmetrically in the DMN but were more weighted towards the contralateral hemisphere in the SMN.

Besides dopamine, dopaminergic neurons co-release GABA via the VMAT2. This hints towards reduced GABA following the VMAT2 knockdown, and consequent imbalance in downstream striatal projection neurons of the direct and indirect pathway [47, 48, 94]. GABA regulates the inhibitory neurotransmission in various brain areas through GABA-A receptors [95]. We hypothesize that decreased binding to the GABA-A receptors may be consistent with loss of inhibitory tone in multiple cortical areas, reflected by our [^11^C]flumazenil PET data, resulting in increased localized brain connectivity, reflected in the fMRI signal. The suppression of the GABAergic feedback circuit, mediated by D2R and external globus pallidus neurons [84], complements the observed elevation in neuronal synchronicity. Our postulation is in line with earlier findings of an inverse correlation of GABA with rs-FC in DMN [96] and with a putative role of loss of inhibitory tone in hyperconnectivity [97]. Interestingly, our [^11^C]flumazenil PET data did not reveal changes in GABA-A expression in the striatum. Instead, we observed a significant decrease in the ipsilateral hemisphere of several cortical regions in both *mild* and *moderate* rats, supporting previous findings of GABA modulation of the internetwork FC [98]. Studies report downregulated inhibitory neurotransmission in Parkinson’s disease, where gene expression of GABAergic markers is low in the frontal cortex [99, 100]. Further, inverse correlations between [^11^C]flumazenil binding and gait disturbances [101], and between GABA concentration in the motor cortex and disease severity have been reported [102]. Moreover, our [^11^C]flumazenil PET results suggest that GABA neurotransmission is disturbed already at the *mild* stage, indicating its potential role as an early biomarker of dopaminergic presynaptic dysfunction. Of note, the observed changes in GABA-A binding might indicate only an apparent decrease in binding due to the reduced GABA availability, or reflect compensatory adaptation on the contralateral hemisphere. Future research should elucidate these aspects and also perform further investigations on glutamate, due to the crucial role of the excitatory/inhibitory-imbalance in several psychiatric disorders [103].

### Limitations and general remarks

We knocked down VMAT2 in dopaminergic projection neurons from the SNc to the dorsal striatum. To achieve selective targeting of this neuronal subtype, rat Cre driver lines have been developed [67, 104]. Although all monoamine-releasing neurons express VMAT2, in contrast to other brain regions, SNc neurons are predominantly dopaminergic [105]. Thus, even though we used wild-type rats, we can largely dismiss effects on other monoaminergic neurons, as also indicated by biochemical analysis of serotonin striatal levels.

Despite its limited off-target editing [38], undesired targeting of SaCas9 on other genes cannot be fully excluded. Nevertheless, off-target candidates with up to 4 mismatches were screened in the whole genome of *Rattus norvegicus* (http://www.rgenome.net/cas-offinder/), consistently with past reports [40]. To the best of our knowledge, the off-target matches (*Ndrg1*, *RGD1305938*, *Btn2a2*, *AABR07042293.2)* have no effects on VMAT2 function, being involved in cell differentiation, T-cell regulation, and mRNA processing, respectively (https://www.ncbi.nlm.nih.gov/IEB/Research/Acembly/index.html).

Another limitation of the study is the relatively small sample size, related to the complex and high-cost procedures involved in the *in vivo* imaging measurements. In addition to this, the intrinsically high intersubject variability in rs-fMRI, and differences in knockdown efficiency contributed to significant variance in our cohorts. Nevertheless, the variability of gene-editing efficiency was in line with previous *in vivo* brain studies [40, 41].

## Conclusions

This work encourages the combinatorial use of CRISPR/Cas9 and molecular and functional *in vivo* brain imaging to achieve selective modulation of genes and understand the related functional adaptations in brain networks, beyond the targeted circuitry.

We anticipate our approach to be a starting point to shed the light on the function of specific genes and their encoded proteins on whole-brain connectivity, useful to understand the cellular basis of functional changes, identify early neurobiological markers, and promising therapeutic interventions.

## Methods

### Animals

Female wild-type Long Evans rats (224 ± 30 g, n = 57) (Charles River Laboratories, Sulzfeld, Germany) were kept on a 12 h day-night cycle at a room temperature of 22 °C and 40 - 60% humidity. Animals received a standard diet and tap water *ad libitum* before and during the experimental period. All animal experiments were performed according to the German Animal Welfare Act and were approved by the local ethical authorities, permit numbers R15/19M, R4/20G.

### Viral vectors

SgRNAs targeting the second exon of the *Slc18a2* gene and the first exon of the *lacZ* gene were designed based on the PAM sequence of SaCas9 (NNGRRT) (Table 1). SgRNAs were cloned into an AAV-PHP.EB expression vector containing a GFP reporter sequence, driven by the CMV promoter, for the identification of transduced neurons. A second AAV-PHP.EB construct was produced to express SaCas9, flanked by two nuclear localization sequences (NLS) to allow its translocation into the nuclei. The vector expressed the nuclease via the CAG promoter and contains three HA-tags to visualize the targeted neurons (Fig. 1a). Cloning of the sgRNAs, plasmid construction, as well as the production of concentrated and purified AAV-PHP.EB vectors, delivered at a concentration of 10^13^ gc/mL, were carried by SignaGen Laboratories (Johns Hopkins University, USA). A European patent application has been filed for the AAV-PHP.EB vectors and is currently pending.

The genomic mutation rate was assessed using a different set of AAVs with AAV2/1 serotype, for a conditional design, kindly provided by Matthias Heidenreich (prev. Zhang lab, Broad Institute of MIT and Harvard, Cambridge, USA).

### Rat primary cortical neuron culture

Primary cortical neurons were obtained from rat embryos of a pregnant Sprague Dawley rat from embryonic day 18 (E18) (Charles River, Sulzfeld, Germany). Embryos were decapitated and quickly removed from the mother rat. Cortical dissection was performed in ice-cold HBSS (100 mL 10 × HBSS, 870 mL dH_2_O, 3.3% 0.3 M HEPES pH 7.3 and 1% pen/strep) (LifeTechnologies, Massachusetts, USA). The obtained tissue was washed three times with 10 mL ice-cold HBSS PhenolRed-free (LifeTechnologies, Massachusetts, USA) and then digested at 37 °C for 20 min in 8 mL HBSS with 2.5% trypsin (LifeTechnologies, Massachusetts, USA). Cortices were washed 3 times with 10 mL HBSS containing 3.7% FBS, and then gently triturated in 2 mL HBSS. For the maintenance, neurons were plated on poly-D-lysine-coated 24 well plates (BD Biosciences, Heidelberg, Germany) or coverslips (Neuvitro Corporation, Vancouver, USA) at a density of 16 × 10^4^/well, and cultured in Neurobasal media supplemented with 2% 1 × B27, 0.25% Glutamax, 0.125% Glutamate and 1% pen/strep (LifeTechnologies, Massachusetts, USA) for four days. Afterward, for the immunofluorescence, AAVs carrying the expression of the vectors for the SaCas9 and sgRNAs (1:1 ratio), were added to the culture medium at 200,000 MOI (AAV-PHP.EB-sgRNA-*lacZ:* 1.7 × 10^12^ gc/mL, AVV-PHP.EB-SaCas9: 1.4 × 10^12^ gc/mL, AAV-PHP.EB-sgRNA-*Slc18a2:* 2.1 × 10^12^ gc/mL, final viral volume 37.5 µL/well). For the Surveyor assay, conditional AAVs for SaCas9, sgRNA-*Slc18a2*, Cre-recombinase (AAV2/1, 1:1:0.5 ratio) were used. Neurons were processed 1-week post-viral treatment (Fig. 1b).

### Surveyor assay

To estimate the VMAT2 knockdown efficiency of the designed sgRNA *in vitro*, we evaluated the presence of genetic deletions in rat primary neurons with the Surveyor assay (Surveyor kit, Integrated DNA Technologies, Coralville, USA). One week following the viral infection, the genomic DNA was extracted using the QuickExtract DNA Extraction solution (Epicentre, Madison, USA), according to the manufactureŕs instructions, and was normalized to 100 ng in dH_2_O. 18 - 25 nt primers were designed 200 - 400 bp away from either side of the SaCas9 target site to amplify the loci of interest by touchdown PCR (oligonucleotides used for PCR are provided in Supplementary Table 8). DNA amplification was performed using 0.5 μL Phusion Polymerase (LifeTechologies, Massachusetts, USA), as previously reported [106]. A single band product was visualized on 1.5% agarose gel, isolated, and purified using QIAquick Spin columns (Qiagen, Hilden, Germany), following the supplier’s protocol. 400 ng of the purified PCR product were mixed with 2 μL Taq DNA polymerase buffer (LifeTechnologies, Massachusetts, USA) to allow the cross-annealing of the mutated and wild-type sequences. The re-annealing process was conducted at the following cycling conditions: 10 min at 95 °C, 95 °C to 85 °C at -2 °C/s, hold 1 min, ramp down to 75 °C at -0.3 °C/s, hold 1 min, and so on until 25 °C temperature was reached. Samples were then stored at 4 °C. This cross-annealing procedure converts the mutations into mismatch duplexes (heteroduplexes), which can be recognized by performing nuclease digestion [107]. We digested the annealed products for 20 min at 42 °C using 2.5 μL MgCl_2_(0.15 M), 1 μL Surveyor nuclease, and 1 μL Surveyor enhancer. Digested products were then resolved on a 2.5% agarose gel, stained with 0.01% SYBR Gold DNA (LifeTechnologies, Massachusetts, USA) in 1% TBE buffer. The size of the occurring bands indicated the location of the mutation (expected DNA fragments sizes are provided in Supplementary Table 8). To quantify the knockdown efficiency, the software ImageJ was used. Peak areas of the bands visualized on agarose gel were selected and the percentages of the transduced neurons acquiring the InDel mutation were calculated using the following formula [106]:

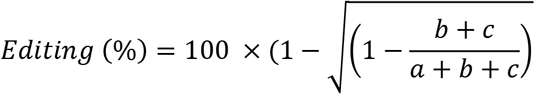

where *a* is the integrated intensity of the undigested PCR product and *b* and *c* are the integrated intensities of each cleavage fragment from the digested product.

### Immunofluorescence of rat primary neurons

Rat primary neurons were processed1-week post-AAV-transduction. Coverslips were washed twice with DPBS and fixed in 4% PFA in DPBS for 15 min at room temperature. Blocking and permeabilization were performed for 30 min in DPBS with 5% donkey serum, 0.5% Triton-X100, and 0.05% BSA. Primary antibody incubation (VMAT2, 1:50, EB06558, Everest Biotech, Ramona, USA, HA-tag: 1:50, 682404, Biolegend, San Diego, USA) was performed for 60 min, followed by Cy3 conjugated secondary antibody incubation (1:250, kindly provided from Birgit Fehrenbacher, Department of Dermatology, University of Tuebingen, Germany). Coverslips were mounted using ProLong Antifade Diamond Medium containing DAPI (LifeTechnologies, Massachusetts, USA), and imaged with a TCS-SP2/Leica DM IRE2 confocal laser scanning microscope. Images were processed with Leica Confocal Software LCS (Version 2.61) (original magnification 630) (Leica Microsystems, Wetzlar, Germany).

### Study design of the *in vivo* experiments

In a first cohort, AAVs-CRISPR/SaCas9 were stereotactically delivered into the right SNc of wild-type rats. Afterward, *in vivo* PET scans with imaging markers of VMAT2 expression, dopamine availability, nerve terminal integrity, and inflammatory responses were performed in VMAT2 knockdown and control rats using [^11^C]DTBZ, [^11^C]raclopride, [^11^C]methylphenidate, and [^18^F]GE- 180, respectively. Motor consequences of the CRISPR/SaCas9-induced VMAT2 knockdown were explored in a wide spectrum of behavioral tasks. Finally, biochemical and histological analyses were performed to corroborate the *in vivo* data (Fig. 2,3,4a).

In a second cohort, cylinder test and [^11^C]raclopride-PET/BOLD-fMRI scans were performed at baseline and after CRISPR/SaCas9-induced VMAT2 knockdown. These measurements were paralleled by *in vivo* PET scans with imaging markers of VMAT2 and GABA-A expression, to inspect the extent of the induced VMAT2 knockdown and its impact on GABA signaling, using [^11^C]DTBZ and [^11^C]flumazenil, respectively (Fig. 5a).

### Stereotactic injections

Rats were anesthetized by injecting a mixture (1 mL/kg) of fentanyl (0.005 mg/kg), midazolam (2 mg/kg,) and medetomidine (0.15 mg/kg) intraperitoneally, and were placed onto a stereotactic frame (Harvard Apparatus, Holliston, MA, USA) with the skull flat between Lambda and Bregma. The following coordinates were used for the injections (flat skull position): AP: - 5 mm, ML: ± 2 mm, DV: - 7.2 mm, below the dural surface as calculated relative to Bregma. Rats were injected with 3 µL AAVs into the right SNc and were divided into two groups. Control rats (n= 10) were injected with AAV-PHP.EB-sgRNA-*lacZ* (1.7 × 10^12^ gc/mL) and AVV-PHP.EB-SaCas9 (1.4 × 10^12^ gc/mL) (1:1 ratio). VMAT2 knockdown rats (n= 47) were injected with AAV-PHP.EB-sgRNA-*Slc18a2* (2.1 × 10^12^ gc/mL) and AVV-PHP.EB-SaCas9 (1.4 × 10^12^ gc/mL) (1:1 ratio) (see Fig. 1a for vector constructs). DPBS (3 µL) was sham-injected into the contralateral SNc. Solutions were infused at a rate of 0.2 µL/min using a 5 µL Hamilton syringe (Hamilton, Bonaduz, Switzerland) and an automated microsyringe pump (Harvard Apparatus, Holliston, MA, USA). To allow for the diffusion of AAVs into the tissue, the needle was left in place for 10 min, and then slowly retracted at 0.2 mm/min. After the surgery, an antidote containing atipamezole (0.75 mg/kg) and flumazenil (0.2 mg/kg) was injected subcutaneously. The rats were kept warm in their cages until fully recovered.

### Radiotracer synthesis

[^11^C]CO_2_ was produced on a medical cyclotron (PETtrace 860, GE Healthcare, Uppsala, Sweden) using the ^14^N(p,α)^11^C route and converted to either [^11^C]MeI (methyl iodide) or [^11^C]MeOTf (methyl triflate) using a Tracerlab FX MeI module (GE Healthcare). A Tracerlab FX M module (GE Healthcare) was used for automated methylation, purification, and formulation of the tracers. [^11^C]Dihydrotetrabenazine (DTBZ), a VMAT2 ligand, was synthesized starting from [^11^C]MeI, which was reacted with 1 mg (+)-9-O-desmethyl-dihydrotetrabenazine (ABX, Radeberg, Germany) in 300 µl DMF in the presence of 7.5 µl 5 M NaOH for 3 min at 40 °C [108]. Afterward, it was purified by HPLC and formulated as a sterile pyrogen-free saline solution. The total synthesis time from the end of the beam was 45 min. The radiochemical purity of the final radiotracer was > 95% as determined by HPLC. Molar activity at the time of injection was 96 ± 37 GBq/μmol.

D-threo-[^11^C]methylphenidate, a dopamine transporter ligand, was synthesized by alkylation of D-threo-N-NPS-ritalinic acid (ABX) using [^11^C]MeI [109]. After acidic deprotection, purification, and formulation, the product was obtained with a 37 ± 6% decay-corrected radiochemical yield (from [^11^C]MeI). The total synthesis time was 55 min, and the radiochemical purity of the final formulated radiotracer was > 95% as determined by HPLC analysis. The molar activity was determined at the time of injection as 58 ± 14 GBq/μmol. [^11^C]Raclopride a D2R ligand, was synthesized by alkylation of S-(+)-O-desmethyl-raclopride (ABX) using [^11^C]MeOTf. After purification and formulation, the product was obtained with a 12 ± 4% decay-corrected radiochemical yield (from [^11^C]methyl triflate). The total synthesis time from the end of the beam was 55 min. The radiochemical purity of the final formulated radiotracer was > 95% as determined by HPLC. The molar activity was determined at the time of injection and was calculated as 88 ± 41 GBq/μmol. [^11^C]Flumazenil, a GABA-A ligand, was synthesized by methylation of desmethylflumazenil (ABX) with [^11^C]MeI. 2 mg of the precursor were dissolved in 0.3 ml DMF with 3 µl 5 M NaOH and reacted for 2 min at 80 °C. After the reaction, the product was purified by semi-preparative HPLC and reformulated by solid-phase extraction (Strata-X, Phenomenex; elution with 0.5 ml ethanol, dilution with 5 ml phosphate-buffered saline). The total synthesis time from the end of the beam was 50 min. The radiochemical purity of the final formulated radiotracer was > 95% as determined by HPLC. The molar activity was determined at the time of injection as 109.5 ± 39.6 GBq/μmol. [^18^F]Fluoride was produced on a PETtrace 860 medical cyclotron (GE Healthcare) using the ^18^O(p,n)^18^F route. [^18^F]GE-180, a translocator protein ligand [110], was synthesized on a FASTlab synthesizer (GE Healthcare) with precursor and reagent kits in single-use cassettes (GE Healthcare) according to the manufacturer’s instructions. The radiochemical purity of the final formulated radiotracer was > 95% as determined by HPLC. The molar activity was determined at the time of injection as 576 ± 283 GBq/μmol.

### *In vivo* PET imaging and data analysis

For the study of cohort 1, VMAT2 knockdown (n= 14) and control rats (n= 10) underwent 60 min dynamic PET emission scans with [^11^C]DTBZ (8 – 10 weeks post-injection), [^11^C]methylphenidate (10 – 12 weeks post-injection), [^11^C]raclopride (12 – 14 weeks post-injection) and [^18^F]GE-180 (14 – 16 weeks post-injection) (Fig. 2a). Four rats were excluded from the data analyses because two rats from each group died during a PET acquisition. One control rat was excluded from the [^11^C]methylphenidate analysis due to a poor signal-to-noise ratio.

For the study of cohort 2, VMAT2 knockdown rats (n= 33), underwent 60 min dynamic PET emission acquisitions with [^11^C]DTBZ (8 – 10 weeks post-injection), and [^11^C]flumazenil (10 – 12 weeks post-injection). The final cohort included 23 rats (see paragraph Simultaneous PET/fMRI experiments).

Three small-animal PET scanners (Inveon, Siemens, Erlangen, Germany) and dedicated rat brain beds (Jomatik Gmbh, Tuebingen, Germany) with stereotactic holders and temperature feedback control units (Medres, Cologne, Germany) were used. These ensured the delivery and removal of the anesthesia gas and stabilized the body temperature at 37 °C during the PET data acquisition. Anesthesia was induced by placing the animals in knock-out boxes and delivering 2% isoflurane in oxygen air. Subsequently, a 24 G catheter (BD Insyte, NJ, USA) was placed into the tail vein for the tracer and/or *i.v.* anesthesia administration. Afterward, animals of cohort 1 were anesthetized with 2% isoflurane vaporized in 1.0 L/min of oxygen. Animals of cohort 2 received a medetomidine bolus injection (0.05 mg/kg) and the anesthesia was switched to constant medetomidine infusion (0.1 mg/kg/h), and 0.5% isoflurane in air during the scan time, as adapted from the literature [111].

The rats were placed in the center of the field of view and PET acquisitions started 5 s before the bolus injection of the tracer. In Supplementary Tables 9 and 10, injected activity (MBq/kg) and molar activity (GBq/µmol) at the time of injection are reported for each radioligand. The list-mode data from the dynamic acquisitions of [^11^C]DTBZ, [^11^C]raclopride, [^11^C]methylphenidate, and [^11^C]flumazenil were histogrammed into 39 time-frames (12×5 s, 6×10 s, 6×30 s, 5×60 s, 10×300 s), from [^18^F]GE-180 into 16 time-frames (5×60 s, 5×120 s, 3×300 s, 3×600 s). PET scans of the study cohort 1 were reconstructed using the OSEM3D map algorithm, and a matrix size of 256 × 256 × 159, resulting in a pixel size of 0.38 × 0.38 × 0.79 mm. PET scans of the study cohort 2 were reconstructed using the OSEM2D algorithm, and a matrix size of 256 × 256 × 89, resulting in a pixel size of 0.33 × 0.33 × 0.79 mm.

Data preprocessing analysis was performed with Matlab (Mathworks, Natick, MA, USA), Statistical Parametric Mapping 12 (SPM12, Wellcome Trust Centre for Neuroimaging, University College London, England), and the QModeling toolbox [112]. First, realignment of all frames was performed using SPM12 and average images were generated for every scan. The mean images were then used for coregistration to the Schiffer rat brain atlas provided by PMOD software [113].

To generate the respective time activity curves (TAC), volumes of interest (VOIs) were defined over the target and reference regions. VOIs were placed over the right and left striatum for [^11^C]DTBZ, [^11^C]raclopride, and [^11^C]methylphenidate, and over the regions reported in Table 2 for [^11^C]flumazenil. Cerebellum was used as reference region for [^11^C]DTBZ, [^11^C]raclopride and [^11^C]methylphenidate. Pons was used as reference region for [^11^C]flumazenil.

Binding potentials for [^11^C]DTBZ, [^11^C]raclopride, [^11^C]methylphenidate and [^11^C]flumazenil were calculated over the all frames in the regions of interest with Logan reference [114], with a population average k2’ ([^11^C]DTBZ: 0.41 min^-1^, [^11^C]raclopride: 0.34 min^-1^, [^11^C]methylphenidate: 0.18 min^-1^, [^11^C]flumazenil: 0.27 min^-1^). [^11^C]DTBZ and [^11^C]raclopride binding changes (%), here expressed as Δ binding, were calculated according to the formula:

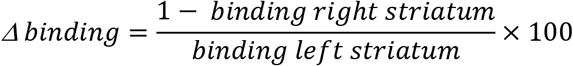

[^18^F]GE-180 uptake in the right striatum was calculated over the interval between 30 – 60 min after scan start and normalized by the uptake calculated in the left (DPBS-injected) striatum over the same time interval.

QModeling was used to generate voxel-wise binding potential maps for [^11^C]DTBZ, [^11^C]raclopride, [^11^C]methylphenidate, and [^11^C]flumazenil. [^18^F]GE-180 average uptake images were generated with an in-house-written script in MATLAB.

### Simultaneous PET/fMRI experiments

Rats (n= 33) underwent longitudinal simultaneous [^11^C]raclopride-PET/BOLD-fMRI scans at baseline and 8 – 14 weeks after CRISPR/SaCas9-induced VMAT2 knockdown. Three rats had to be excluded from the data analyses due to aliasing artifacts, local distortion, and motion during the data acquisition. Seven rats died during a PET/BOLD-fMRI scan. The final cohort included 23 rats. Anesthesia induction and injection were performed as described above for the cohort 2 (see paragraph *In vivo* PET imaging and data analysis).

Next, rats were placed onto a dedicated rat bed (Medres, Cologne, Germany) and a temperature feedback control unit (Medres, Cologne, Germany), ensuring the delivery and removal of the anesthesia gas and stabilizing the body temperature at 37 °C during the scan time. A breathing pad and a pulse oximeter were used to observe respiration and heart rates. Scans were acquired using a small-animal 7 T Clinscan MRI scanner, a 72 cm diameter linearly polarized RF coil (Bruker) for transmission, and a four-channel rat brain coil for reception (Bruker Biospin MRI, Ettlingen, Germany). Localizer scans were first acquired to accurately position the rat brains into the center of the PET/MRI field of view. Subsequently, local field homogeneity was optimized by measuring local magnetic field maps. Anatomical reference scans were performed using T2- weighted Turbo-RARE MRI sequences (TR: 1800 ms, TE: 67.11 ms, FOV: 40 x 32 x 32 mm, image size: 160 x 128 x 128 px, Rare factor: 28, averages: 1). Finally, T2*-weighted gradient echo EPI sequences (TR: 2500 ms, TE: 18 ms, 0.25 mm isotropic resolution, FoV 25 x 23 mm, image size: 92 x 85 x 20 px, slice thickness 0.8 mm, slice separation 0.2 mm, 20 slices) were acquired for BOLD-fMRI.

A dedicated small-animal PET insert developed in cooperation with Bruker (Bruker Biospin MRI, Ettlingen, Germany) was used for the [^11^C]raclopride acquisitions, the second generation of a PET insert developed in-house with similar technical specifications [115]. [^11^C]Raclopride was applied via a bolus injection. In Supplementary Table 10 injected activities (MBq/kg) and molar activities (GBq/µmol) at the time of injection are reported. PET/fMRI acquisitions started simultaneously with the tracer injection and were performed over a period of 60 min. The list-mode files of the PET data were histogrammed into 14 time-frames (1x30 s, 5x60 s, 5x300 s, 3x600 s), the 30s between acquisition start and the injection were excluded from the analysis. Reconstruction was performed with an in-house-written OSEM2D algorithm. Data preprocessing and analysis were performed as described above (see paragraph *In vivo* PET imaging and data analysis). A population average k2’ ([^11^C]raclopride baseline: 0.20 min^-1^, [^11^C]raclopride VMAT2 knockdown: 0.23 min^-1^) was set for the Logan reference [114].

Preprocessing of the fMRI data was performed using a pipeline employing SPM12, Analysis of Functional NeuroImages (AFNI, National Institute of Mental Health (NIMH), Bethesda, Maryland, USA), and in-house-written scripts in MATLAB, as reported previously [116].

RS-FC was calculated using a seed-based approach. To this extent, 20 regions were selected from the Schiffer rat brain atlas (a list of the regions is provided in Table 2). The SPM toolbox Marseille Boîte À Région d’Intérêt (MarsBaR) was employed to extract fMRI time-courses from all regions [117]. These were then used to calculate pairwise Pearson’s r correlation coefficients for each dataset, generating correlation matrices containing 20 x 20 elements. Self-correlations were set to zero. The computed Pearson’s r coefficients then underwent Fischer’s transformation into z values for group-level analysis.

Several rs-FC metrics were computed on different regional levels to investigate the potential effects of dopamine depletion in the right striatum. Regional node strengths were computed as the sum of all correlations of one seed to the regions included in one network. Interregional node strengths were defined as the sum of the correlations of one node to the regions of another network. Network strengths were defined as the sum of strengths of all correlations between regions belonging to a network. Internetwork strengths were calculated as the sum of all correlations between two sets of regions belonging to two networks [118].

### Behavioral analysis

#### Cylinder test

Untrained rats were placed individually inside a glass cylinder (19 cm Ø, 20 cm height). The test started immediately and lasted 5 min. During the test session rats were left undisturbed and were videotaped with a camera located at the bottom-center of the cylinder to allow a 360 ° angle view.

Paw touches were analyzed using a slow-motion video player (VLC software, VideoLan). The number of wall touches, contacts with fully extended digits, was counted. Data were analyzed as follows:

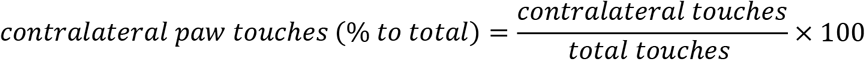

One rat from the cohort 2 was excluded from the analysis due to issues during video recording.

#### Rotameter test

Rotational asymmetry was assessed using an automated rotameter system composed of four hemispheres (TSE Systems GmBH, Bad Homburg, Germany) based on the design of Ungerstedt and Arbuthnott [119]. Rats were placed into an opaque half bowl (49 cm Ø; 44 cm height) and fixed to a moveable wire with a collar. The number of clockwise (CW) and counterclockwise (CCW) rotations (difference of 42.3° in their position) were automatically counted. The spontaneous rotation test lasted 5 min. Apomorphine-evoked rotational asymmetry was evaluated for 60 min after *s.c*. administration of apomorphine hydrochloride (0.25 mg/kg) dissolved in physiological saline containing 0.1% ascorbic acid (Sigma Aldrich, St. Louis, Missouri, USA). Two priming injections of apomorphine (1-week interval off-drug) were necessary to produce sensitization to the treatment. The program RotaMeter (TSE Systems GmBH, Bad Homburg, Germany) was used to acquire the data. Data were analyzed as follows:

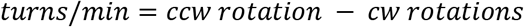

#### Beam walk test

Rats were trained for 4 days to cross a built-in-house beam (1.7 cm width, 60 cm length, 40 cm height), and reach a cage with environmental enrichment. Rats had 5 trials to cross the beam with the reward resting time decreasing from 30 s to 10 s. On the test day (one week apart from the fourth day of training), rats were videotaped. The acquired videos were analyzed using a slow-motion video player (VLC software, VideoLan) and the number of footslips (falls) was counted. Data were analyzed as follows:

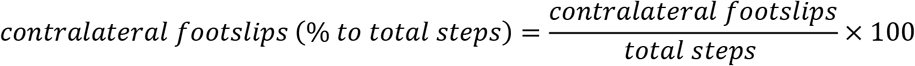

#### Open Field test

Untrained rats were set in a rectangular box (TSE Systems GmBH, Bad Homburg, Germany) for 11 min (1 min habituation, 10 min test) to evaluate the spontaneous exploratory behavior. A frame with light sensors, as high as the animals’ body center, was connected to a receiver box to record the rats’ walked distance. The program ActiMod (TSE Systems GmBH, Bad Homburg, Germany) was used for the analysis and the experimental session was divided into 1 min time bins. Results were averaged from the total traveled distance over 10 min (mean ± SD).

#### Body weight gain

Rats’ body weight was measured before (week 0) and 14 weeks after CRISPR/SaCas9-induced VMAT2 knockdown. Body weight gain was calculated as follows:

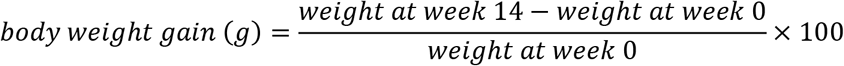

### Histology of rat brain slices

Rats of cohort 1 (n= 20) were sacrificed 19 weeks after viral vector injection via CO_2_inhalation followed by intracardial perfusion with heparinized DPBS (1:50 v/v, 100 mL). After decapitation, brains were rapidly removed and placed in a brain matrix on ice. The left and right striata were dissected, from 2 mm thick coronal sections, flash-frozen in liquid nitrogen, and stored at – 80 °C until further analysis (HPLC). The remaining tissue was fixed in 4% paraformaldehyde for 48 h and then transferred to a 20% sucrose solution for cryoprotection. Brains were cut into 35 µm thick coronal sections on a freezing microtome (Leica Biosystems, Wetzlar, Germany) and stored in an anti-freeze solution (0.5 M phosphate buffer, 30 % glycerol, 30 % ethylene glycol) at - 20 °C. Sections were collected in 12 equally spaced series through the entire anterior-posterior extent of the SN and striatum and stored until further analysis. Immunohistochemistry was performed on free-floating sections. Sections were washed 3 times with TBS buffer and antigen retrieval was carried for 30 min at 80 °C in Tris/EDTA buffer. Afterward, pre-incubation in MeOH (10%) and H_2_O_2_(3%) in TBS was performed for 30 min. Following the blocking in 5% normal goat serum in TBS-X (0.05%), primary antibody incubation was performed for 24 h at room temperature in 1% BSA in TBS-X (TH: 1:5000, P40101, Pel-freez, Arkansas, USA, VMAT2: 1:5000, 20042, Immunostar, Hudson, USA). The tissue was rinsed in TBS-X and reacted with the respective biotinylated secondary antibody (1:200, Vector Laboratories Ltd., Peterborough, UK) for 60 min at room temperature in 1% BSA in TBS-X. Staining was developed using 3,3’- diaminobenzidine (DAB Substrate Kit, Vector Laboratories Ltd., Peterborough, UK) and an immunoperoxidase system (Vectastain Elite ABC-Kit, Vector Laboratories Ltd., Peterborough, UK). Slices were rinsed, mounted onto chromalum gelatinized slides, dehydrated in ascending concentrations of alcohol and xylene baths, and coverslipped with DPX mounting medium (Sigma Aldrich, St. Louis, Missouri, USA).

### Stereological analysis

Estimates of total numbers of TH+ cells in nigral sections were obtained with an unbiased stereological quantification method by employing the optical fractionator principle [120]. Brain sections from 5 rats (VMAT2 knockdown n= 2, Control n= 3) were excluded from the stereological analysis, due to weak TH immunoreactivity. First, 5x images were acquired with the automated Metafer slide scanning platform (MetaSystems, Altlußheim, Germany). Then, ROIs were drawn using the VSViewer program (Metasystems, Altlußheim, Germany), and a sampling fraction of 50% was defined. Afterward, 63x images were acquired in an automated fashion based on the sampling fraction in a random orientation within the ROI. The acquired 63x images were imported into the VIS program and cell counting was performed with the CAST module (Visiopharm A/S, Hørsholm, Denmark, Version 2020.08.2.8800).

The number of cells estimates was obtained by applying the formula:

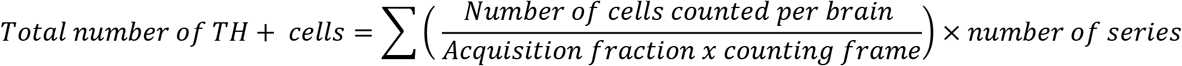

### Immunofluorescence of rat brain slices

Immunofluorescence was performed on SN sections from rats of cohort 1 (35 µm thick coronal sections), mounted onto chromalum gelatinized slides, dehydrated in ascending concentrations of alcohol and xylene baths. Sections were washed 3 times with KPBS buffer and antigen retrieval was carried for 30 min at 80 °C in Tris/EDTA buffer. Following blocking in 5% donkey serum and normal horse serum in KPBS-X (0.25%), primary antibody incubation was performed for 24 h at room temperature (VMAT2: 1:5000, 20042, Immunostar, Hudson, USA, HA-tag: 1:5000, MMS- 101R, Nordic BioSite, Taby, Sweden, GFP: 1:50.000, ab13970, Abcam, Cambridge, UK). The tissue was rinsed in KPBS-X and reacted with the respective fluorophore-conjugated secondary antibody (1:200, Vector Laboratories Ltd., Peterborough, UK) for 2 h in 0.2% KPBS-X. Slides were coverslipped with Vectashield mounting medium (Vector Laboratories Ltd., Peterborough, UK).

### Biochemistry

Dopamine, 3,4-Dihydroxyphenylacetic acid (DOPAC), homovanillic acid (HVA), and serotonin (5- HT) striatal levels were determined by HPLC. Brain samples from 5 rats (VMAT2 knockdown n= 3, Control n= 2) were excluded from the analysis as striatal sections did not fall in the selected range: 1.2 – 0.4 mm.

Briefly, striatal brain lysates generated from cohort 1 were injected by a cooled autosampler into an ESA Coulchem III coupled to a Decade Elite electrochemical detector (Antec Scientific, Zoeterwoude, The Netherlands) set to a potential of +350 mV. Separation was facilitated by using an Atlantis Premier BEH C18 AX column (Waters Corporation, Massachusetts, USA) and a dual mobile phase gradient of decreasing octane sulfonic acid (OSA) and increasing MeOH content (mobile phase A containing 100 mM PO_4_-buffer pH 2.50 and 4.62 mM OSA and mobile phase B containing 100 mM PO_4_-buffer pH 2.50 and 2.31 mM OSA), delivered at a flow rate of 0.35 mL/min to an Atlantis Premier BEH C18 column (particle size 2.5 µm, 2.1 mm x 150 mm) (Waters Corporation, Massachusetts, USA). Data was collected using the Chemstation software (Agilent, California, USA) and then exported to Chromeleon (LifeTechnologies, Massachusetts, USA) for data quality control, peak integration, and concentration calculations. Striatal metabolites’ content was expressed in nmol for each sample and normalized to total protein (mg). Dopamine turnover rate was calculated according to the formula:

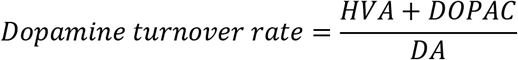

### Statistics

Statistical analysis was performed with GraphPad Prism 9.0 (Graphpad Software) if not otherwise stated. Results were analyzed using paired t-tests for the within-subjects comparisons, and unpaired t-tests for the between-groups comparisons. Correlations were performed using linear regression analyses. Synaptic dysfunction discrimination was tested with multiple-comparison ANOVA.

## Acknowledgments

We are very grateful to Feng Zhang and all members of the laboratory, Broad Institute of MIT and Harvard, for the help and support with establishing the CRISPR gene knockdown experiments. We thank the technical assistants Sandro Aidone, Daniel Bukala, Linda Schramm, Maren Harant, Ramona Stremme, Elena Kimmerle and Johannes Kinzler. We also thank Ulla Samuelsson, Ulrika Sparrhult-Björk, Dr. Ulrika Schagerlöf, Anneli Josefsson and Anna Hansen at Lund University, Sweden for their technical support. Additionally, we acknowledge Dr. Julia Mannheim, Dr. Andreas Schmid, Dr. Rebecca Rock, Ines Herbon, Dr. Neele Hübner, Dr. Andreas Dieterich, Hans Jörg Rahm, Dr. Carsten Calaminus, Funda Cay, Kristin Patzwaldt, Laura Kübler, Marilena Poxleitner, Sabrina Buss and Dominik Seyfried for their administrative, technical, and experimental support at the Department of Preclinical Imaging and Radiopharmacy, Werner Siemens Imaging Center, Eberhard Karls University, Tuebingen. This study is also part of the PhD theses of Sabina Marciano and Tudor Mihai Ionescu.

## Funding

• German Research Foundation to KH

• Carl-Zeiss Foundation to KH

• Werner Siemens Foundation to BJP

• Deutscher Akademischer Austauschdienst to SM, KH

## Author contributions

• Conceptualization: KH

• Methodology: SM, TMI

• Software: SM, TMI

• Validation: SM

• Formal analysis: SM, TMI

• Investigation: SM, TMI, AM, RSS, RYC

• Resources: BJP, DK

• Data curation: SM

• Writing – original draft: SM

• Writing – review and editing: SM, TMI, AM, DK, RSS, RYC, BJP, KH

• Supervision: KH

• Project administration: KH

• Funding acquisition: BJP, KH

## Data availability

The original dataset will be made available upon request.

## Supplementary Figures

**Supplementary Fig. 1.**
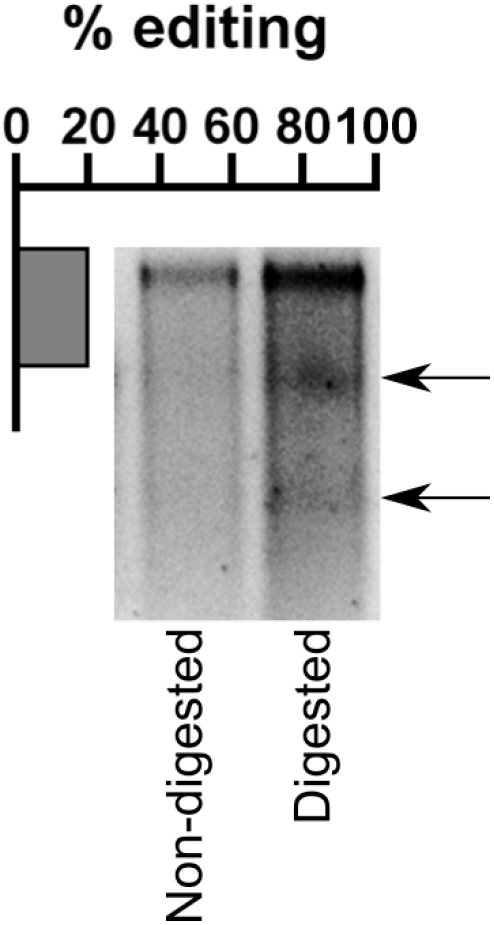
*In vitro* validation of CRISPR/SaCas9-induced VMAT2 knockdown in rat primary cortical neurons. One-week post-transduction with conditional vectors for SaCas9 and sgRNA- *Slc18a2*, neurons were processed for the Surveyor assay. The expected cleavage products (arrows), and estimated editing of 20%, could be seen for the digested DNA.

**Supplementary Fig. 2.**
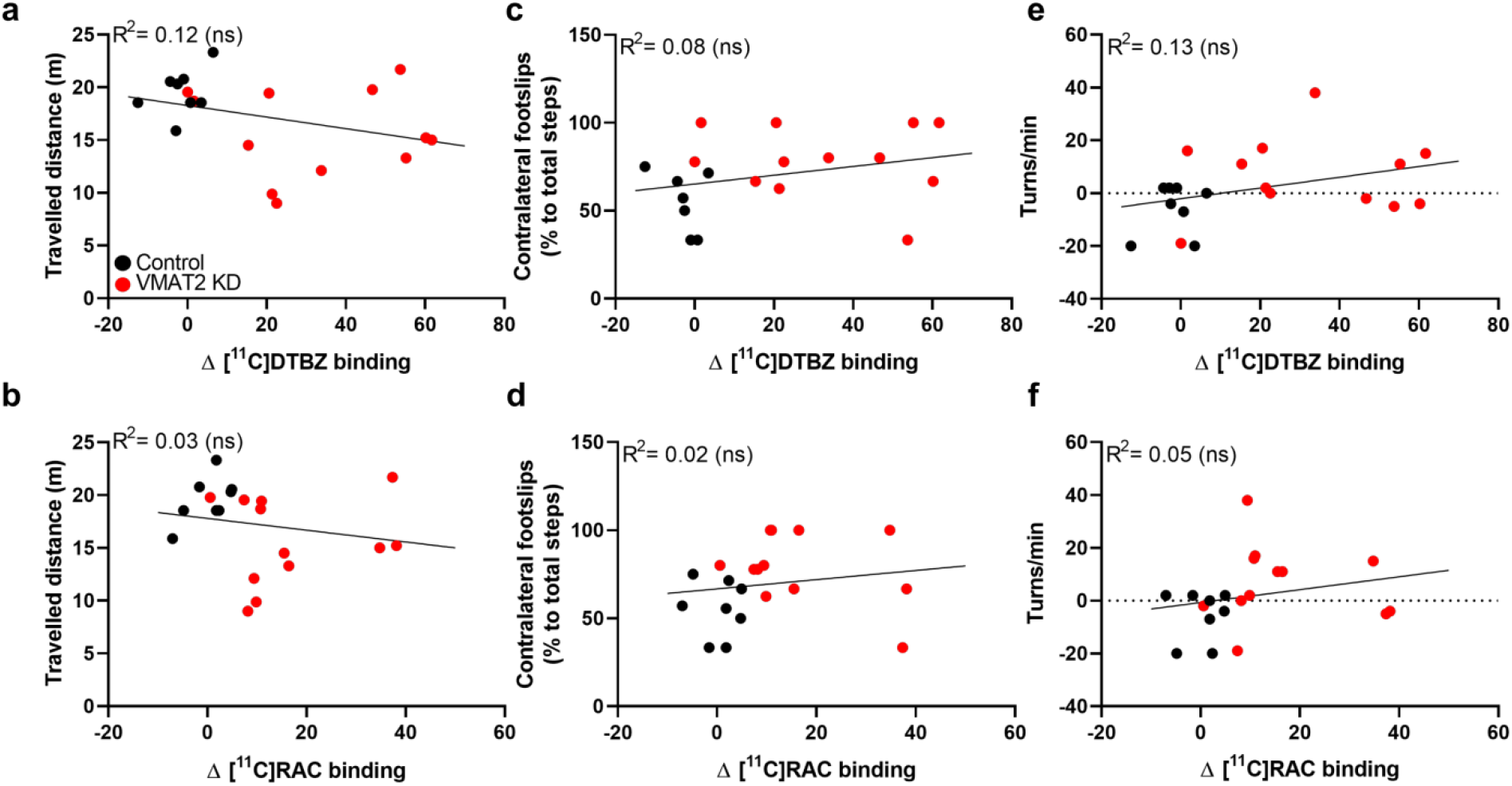
CRISPR/SaCas9-induced VMAT2 knockdown, postsynaptic changes, and motor impairment do not correlate for whole-body movements. No correlation was observed between locomotor activity and (**a**) VMAT2 expression changes (Δ [^11^C]DTBZ binding), or (**b**) dopamine availability (Δ [^11^C]RAC binding). The number of contralateral footslips in the beam walk test did not correlate with (**c**) changes in VMAT2 expression (Δ [^11^C]DTBZ binding) or (**d**) dopamine availability (Δ [^11^C]RAC binding). Spontaneous rotations did not correlate with (**e**) changes in VMAT2 expression (Δ [^11^C]DTBZ binding) or (**f**) dopamine availability (Δ [^11^C]RAC binding). ns *P*≥ 0.05. Control rats n= 8; VMAT2 KD rats n= 12. KD, knockdown; [^11^C]RAC, [^11^C]raclopride.

**Supplementary Fig. 3.**
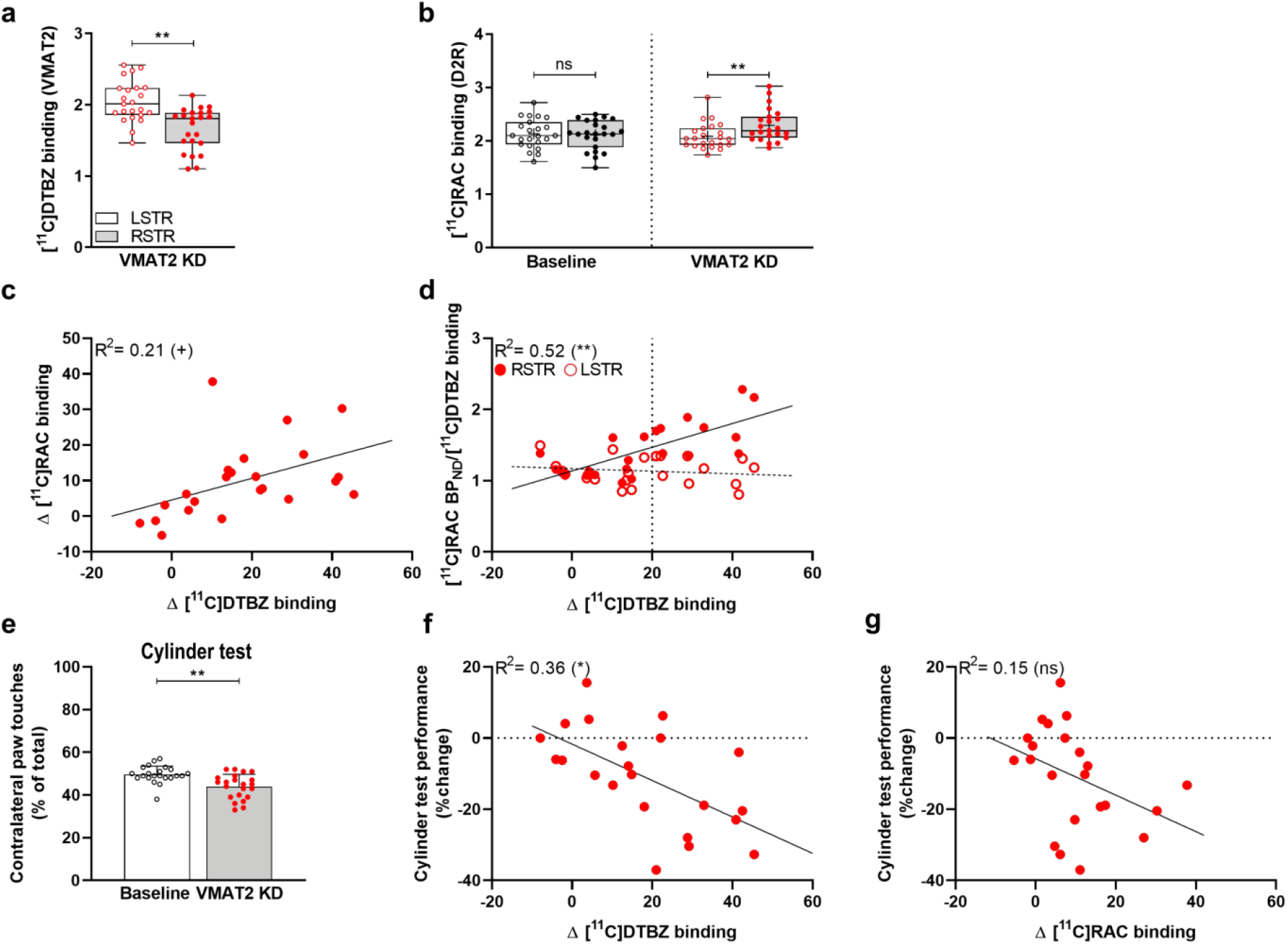
Reproducibility of the CRISPR/SaCas9-induced VMAT2 knockdown. (**a**) [^11^C]DTBZ in VMAT2 KD rats and (**b**) [^11^C]RAC PET at baseline and after CRISPR/SaCas9-induced VMAT2 KD. (**c**) A strong correlation between Δ [^11^C]RAC and Δ [^11^C]DTBZ binding is shown. (**d**) At 20% VMAT2 KD (Δ [^11^C]DTBZ binding), D2R binding (Δ [^11^C]RAC binding) prominently increased. This threshold was therefore set to separate the rats into *mild* and *moderate*. Data are shown as boxplot with the median value (central mark), the mean value (plus sign), interquartile range (boxes edges), and the extreme points of the distribution (whiskers). Baseline n= 23, VMAT2 KD n= 23. (**e**) Cylinder test at baseline (n= 22) and 12 - 14 weeks after CRISPR/SaCas9 gene-editing (n= 22). VMAT2 KD rats showed reduced contralateral paw touches compared to baseline. Rats performance in the cylinder test, % change from baseline, correlated with reduced VMAT2 expression (Δ [^11^C]DTBZ binding) (**f**), but not with dopamine availability (Δ [^11^C]RAC binding) (**g**). Data are shown as mean ± SD. *^+^P*< 0.05, *\*P*< 0.01, *\*\*P*< 0.001. KD, knockdown; [^11^C]RAC, [^11^C]raclopride.

**Supplementary Fig. 4.**
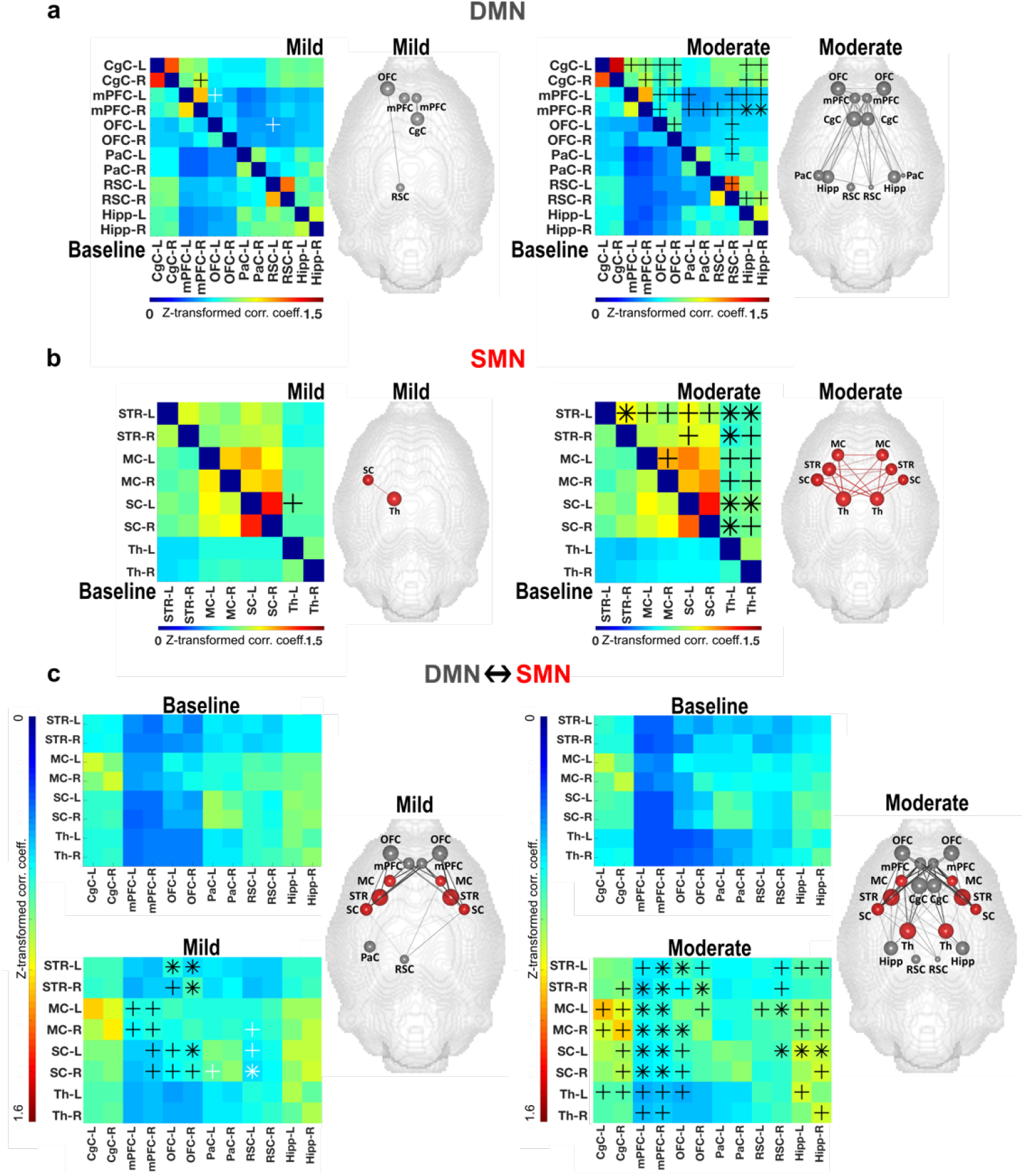
Increased resting-state functional connectivity after CRISPR/SaCas9-induced VMAT2 knockdown. Group level correlation matrices of the DMN (**a**) and SMN (**b**) at baseline and after VMAT2 KD for *mild* (left panel) and *moderate* (right panel) rats. Brain graphs, right to the matrices, illustrate the nodes and edges (raw values) that demonstrated rs-FC changes to baseline (%) in DMN (**a**) and SMN (**b**). (**a**) In rats with *mild* VMAT2 KD, we observed divergent rs-FC changes to baseline in regions of the anterior DMN (left mPFC and OFC, right mPFC and CgC). In rats with *moderate* VMAT2 KD, we observed rs-FC increase in most regions of the anterior DMN, extending to the posterior DMN (RSC bilaterally, right RSC, and Hipp bilaterally). (**b**) In rats with *mild* VMAT2 KD, we observed rs-FC increase between the contralateral SC and Th, which intensified in *moderate* VMAT2 KD rats and extended to most of the regions of the SMN. (**c**) Group level correlation matrices of the DMN-SMN rs-FC changes at baseline and after VMAT2 KD for *mild* (left panel) and *moderate* (right panel) rats. Brain graphs, right to the matrices, illustrate the nodes and edges (raw values) that demonstrated internetwork rs-FC changes to baseline (%). Internetwork FC changes in rats with *mild* VMAT2 KD indicated increased rs-FC between anterior regions of the DMN and the SMN (between OFC and STR, SC bilaterally, between right mPFC and MC, SC bilaterally), and decreased rs-FC between posterior regions of the DMN and the SMN (between left RSC and right MC, and SC bilaterally). Increased DMN-SMN rs-FC was found in rats with *moderate* VMAT2 KD. Internetwork rs-FC alterations extended between CgC/Hipp, and SC, MC, Th. *^+^P*< 0.05, *\*P*< 0.01. *Mild*: Δ [^11^C]DTBZ binding < 20%, n= 13; *Moderate*: Δ [^11^C]DTBZ binding ≥ 20%, n= 10. KD, knockdown; DMN, default-mode network; SMN, sensorimotor network. mPFC, medial prefrontal cortex; OFC, orbitofrontal cortex; CgC, cingulate cortex; RSC, retrosplenial cortex; Hipp, hippocampus; SC, somatosensory cortex; Th, thalamus; STR, striatum; MC, motor cortex. Abbreviations of brain regions considered for the analysis of the fMRI data, including their respective volumes, are reported in Table 2.

**Supplementary Fig. 5.**
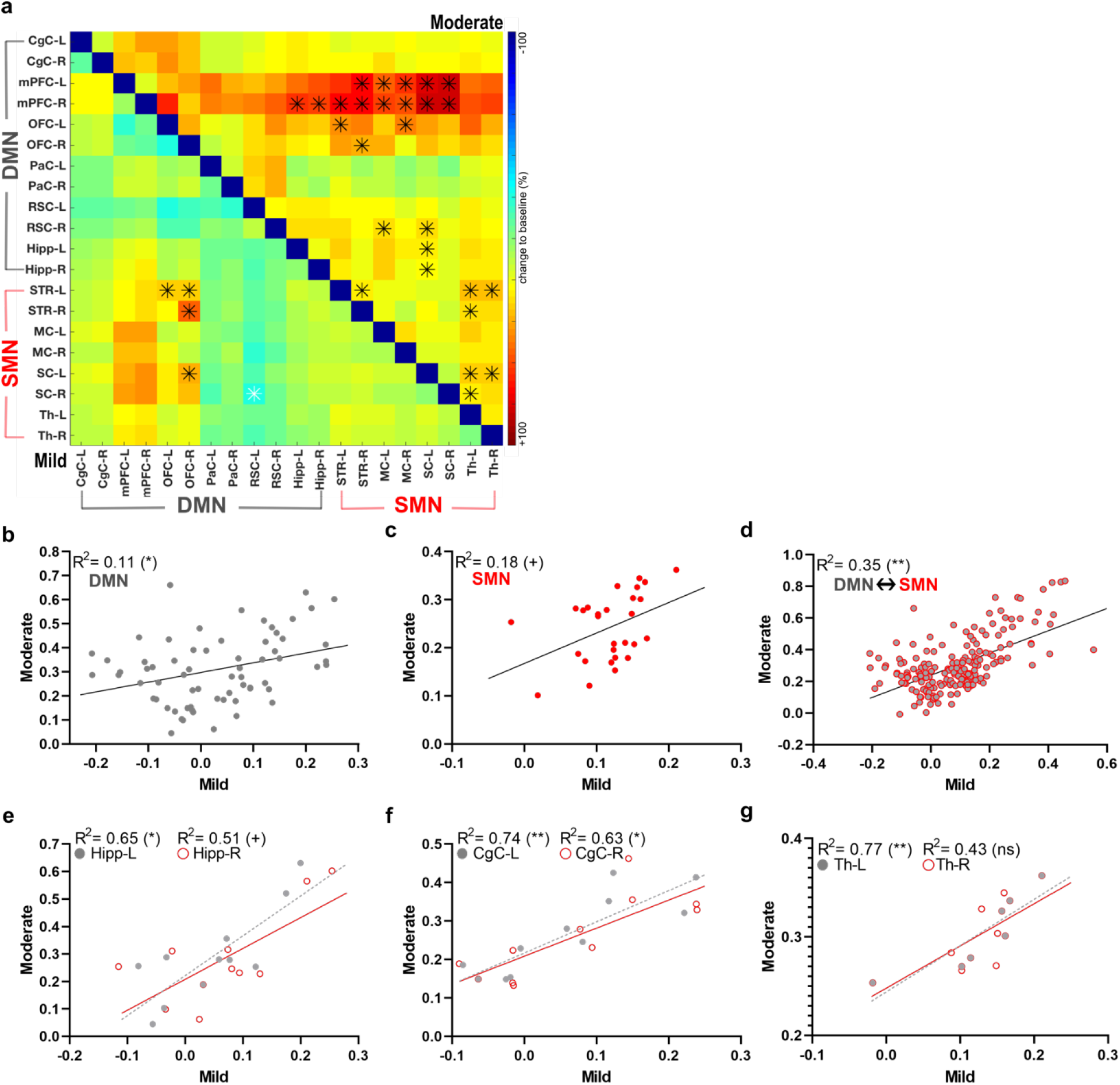
Node correlation analyses evidence pattern similarity and a linear relationship between the magnitude of the resting-state functional connectivity changes in *mild* and *moderate* rats. (**a**) Group level correlation matrices of the intraregional and internetwork rs-FC changes to baseline (%) for *mild* and *moderate* VMAT2 KD rats. (**b-d**) Group level intraregional and internetwork rs-FC changes correlated linearly between the two groups. Node correlation analysis between *mild* and *moderate* VMAT2 KD rats indicated a linear increase in the magnitude of the rs-FC changes to baseline in the right and left Hipp (**e**), and CgC (**f**). (**g**) Rs-FC changes (%) in the SMN correlated linearly between *mild* and *moderate* VMAT2 KD rats in the left, but not in the right Th. Our data suggest a similarity in the pattern of the intraregional and internetwork rs-FC changes between rats with *mild* and *moderate* VMAT2 KD and a linear relationship between the magnitude of the rs-FC changes. ^+^*P*< 0.05, \**P*< 0.01, \*\**P*< 0.001. *Mild*: Δ [^11^C]DTBZ binding < 20%, n= 13; *Moderate*: Δ [^11^C]DTBZ binding ≥ 20%, n= 10. KD, knockdown; DMN, default-mode network; SMN, sensorimotor network. Hipp, hippocampus; CgC, cingulate cortex. Th, thalamus. Abbreviations of brain regions considered for the analysis of the fMRI data, including their respective volumes, are reported in Table 2.

**Supplementary Fig. 6.**
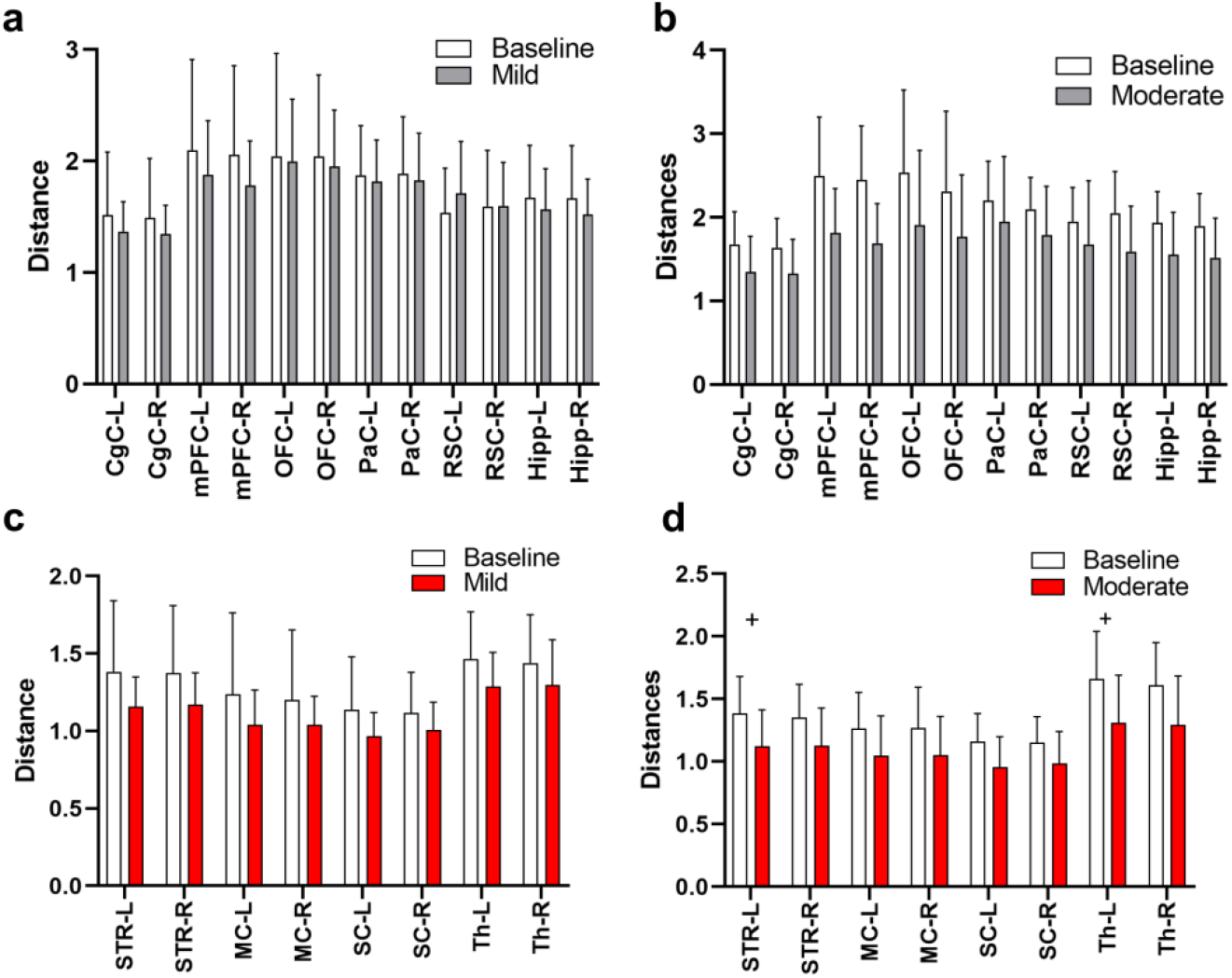
Graph theoretical analyses reveal lateralized recruitment of SMN regions following the CRISPR/SaCas9-induced VMAT2 *moderate* knockdown. Quantitative graph theoretical analyses of the global mean connection distance indicated no changes for *mild* (**a**) and *moderate* (**b**) VMAT2 KD rats in the DMN. (**c**) Rats with *mild* VMAT2 KD displayed no changes in the SMN. (**d**) Instead, rats with *moderate* VMAT2 KD presented shorter functional paths to the nodes in the contralateral Th and STR. Data are shown as mean ± SD. *^+^P*< 0.05, Bonferroni-Sidak-corrected. *Mild*: Δ [^11^C]DTBZ binding < 20%, n= 13; *Moderate*: Δ [^11^C]DTBZ binding ≥ 20%, n= 10. KD, knockdown; DMN: default-mode network, SMN: sensorimotor network; Th: Thalamus; STR: striatum. Abbreviations of brain regions considered for the analysis of the fMRI data, including their respective volumes, are reported in Table 2.

## Supplementary Tables

**Supplementary Table 1:** *P* values of the functional connectivity changes in regions of the DMN for rats with *mild* VMAT2 knockdown. Data were analyzed using paired t-tests.

| *Mild* | Hipp-R | Hipp-L | RSC-R | RSC-L | PaC-R | PaC-L | OFC-R | OFC-L | mPFC-R | mPFC-L | CgC-R |
| --- | --- | --- | --- | --- | --- | --- | --- | --- | --- | --- | --- |
| CgC-L | 0.40 | 0.56 | 0.95 | 0.33 | 0.76 | 0.84 | 0.22 | 0.27 | 0.07 | 0.07 | 0.61 |
| CgC-R | 0.30 | 0.41 | 0.83 | 0.29 | 0.84 | 0.86 | 0.17 | 0.23 | 0.03 | 0.08 |  |
| mPFC-L | 0.21 | 0.34 | 0.77 | 0.73 | 0.36 | 0.55 | 0.46 | 0.02 | 0.53 |  |  |
| mPFC-R | 0.12 | 0.23 | 0.54 | 0.99 | 0.24 | 0.46 | 0.24 | 0.61 |  |  |  |
| OFC-L | 0.49 | 0.55 | 0.20 | 0.05 | 0.51 | 0.67 | 0.18 |  |  |  |  |
| OFC-R | 0.31 | 0.20 | 0.26 | 0.13 | 0.23 | 0.45 |  |  |  |  |  |
| PaC-L | 0.61 | 0.39 | 0.84 | 0.08 | 0.31 |  |  |  |  |  |  |
| PaC-R | 0.74 | 0.59 | 0.83 | 0.11 |  |  |  |  |  |  |  |
| RSC-L | 0.18 | 0.22 | 0.31 |  |  |  |  |  |  |  |  |
| RSC-R | 0.75 | 0.65 |  |  |  |  |  |  |  |  |  |
| Hipp-L | 0.67 |  |  |  |  |  |  |  |  |  |  |

**Supplementary Table 2:**
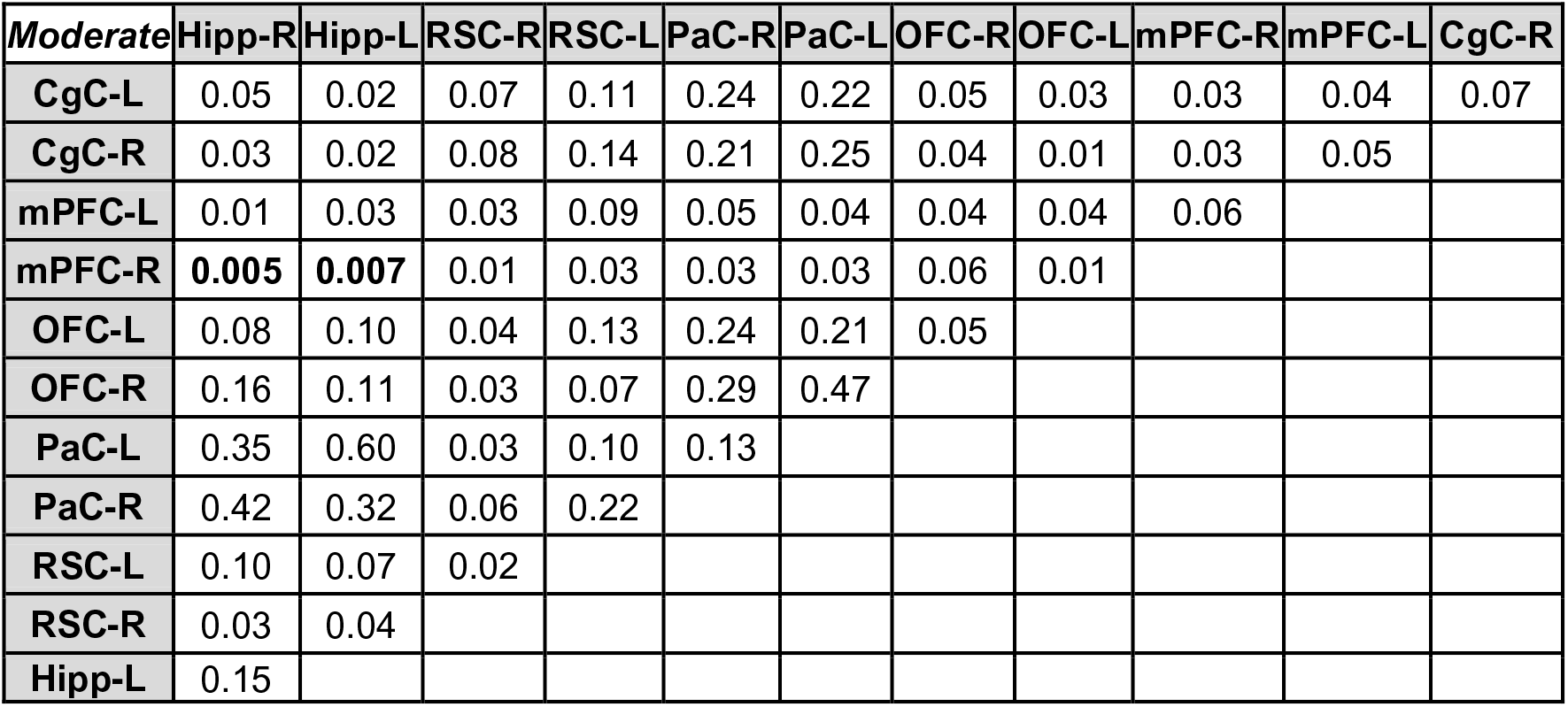
*P* values of the functional connectivity changes in regions of the DMN for rats with *moderate* VMAT2 knockdown. Data were analyzed using paired t-tests.

| *Moderate* | Hipp-R | Hipp-L | RSC-R | RSC-L | PaC-R | PaC-L | OFC-R | OFC-L | mPFC-R | mPFC-L | CgC-R |
| --- | --- | --- | --- | --- | --- | --- | --- | --- | --- | --- | --- |
| CgC-L | 0.05 | 0.02 | 0.07 | 0.11 | 0.24 | 0.22 | 0.05 | 0.03 | 0.03 | 0.04 | 0.07 |
| CgC-R | 0.03 | 0.02 | 0.08 | 0.14 | 0.21 | 0.25 | 0.04 | 0.01 | 0.03 | 0.05 |  |
| mPFC-L | 0.01 | 0.03 | 0.03 | 0.09 | 0.05 | 0.04 | 0.04 | 0.04 | 0.06 |  |  |
| mPFC-R | **0.005** | **0.007** | 0.01 | 0.03 | 0.03 | 0.03 | 0.06 | 0.01 |  |  |  |
| OFC-L | 0.08 | 0.10 | 0.04 | 0.13 | 0.24 | 0.21 | 0.05 |  |  |  |  |
| OFC-R | 0.16 | 0.11 | 0.03 | 0.07 | 0.29 | 0.47 |  |  |  |  |  |
| PaC-L | 0.35 | 0.60 | 0.03 | 0.10 | 0.13 |  |  |  |  |  |  |
| PaC-R | 0.42 | 0.32 | 0.06 | 0.22 |  |  |  |  |  |  |  |
| RSC-L | 0.10 | 0.07 | 0.02 |  |  |  |  |  |  |  |  |
| RSC-R | 0.03 | 0.04 |  |  |  |  |  |  |  |  |  |
| Hipp-L | 0.15 |  |  |  |  |  |  |  |  |  |  |

**Supplementary Table 3:**
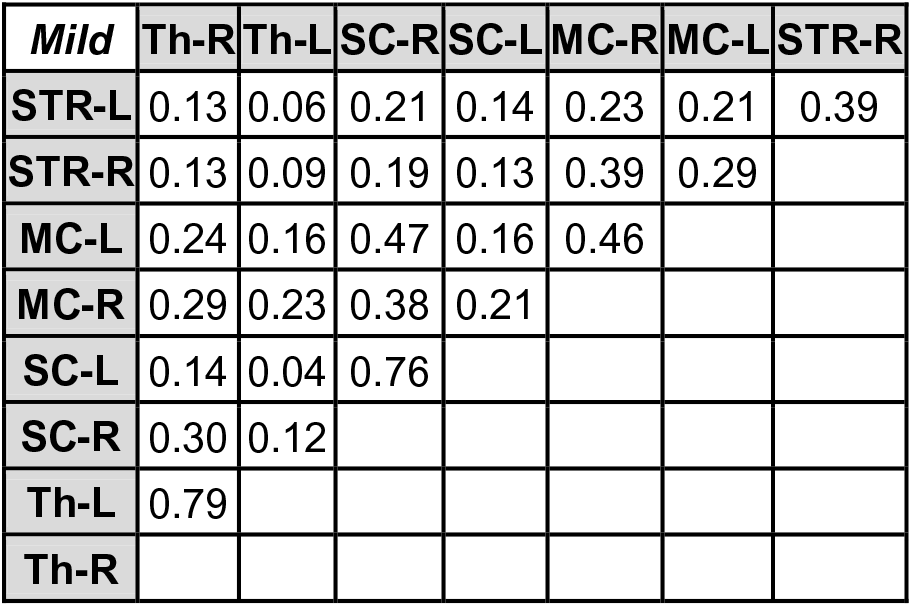
*P* values of the functional connectivity changes in regions of the SMN for rats with *mild* VMAT2 knockdown. Data were analyzed using paired t-tests.

**Supplementary Table 4:**
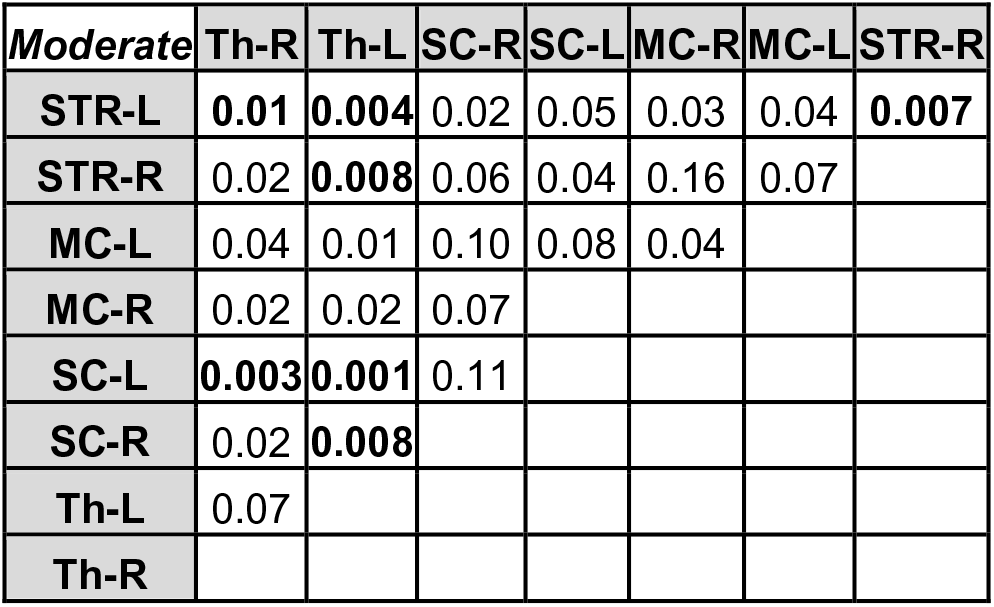
*P* values of the functional connectivity changes in regions of the SMN for rats with *moderate* VMAT2 knockdown. Data were analyzed using paired t-tests.

**Supplementary Table 5:**
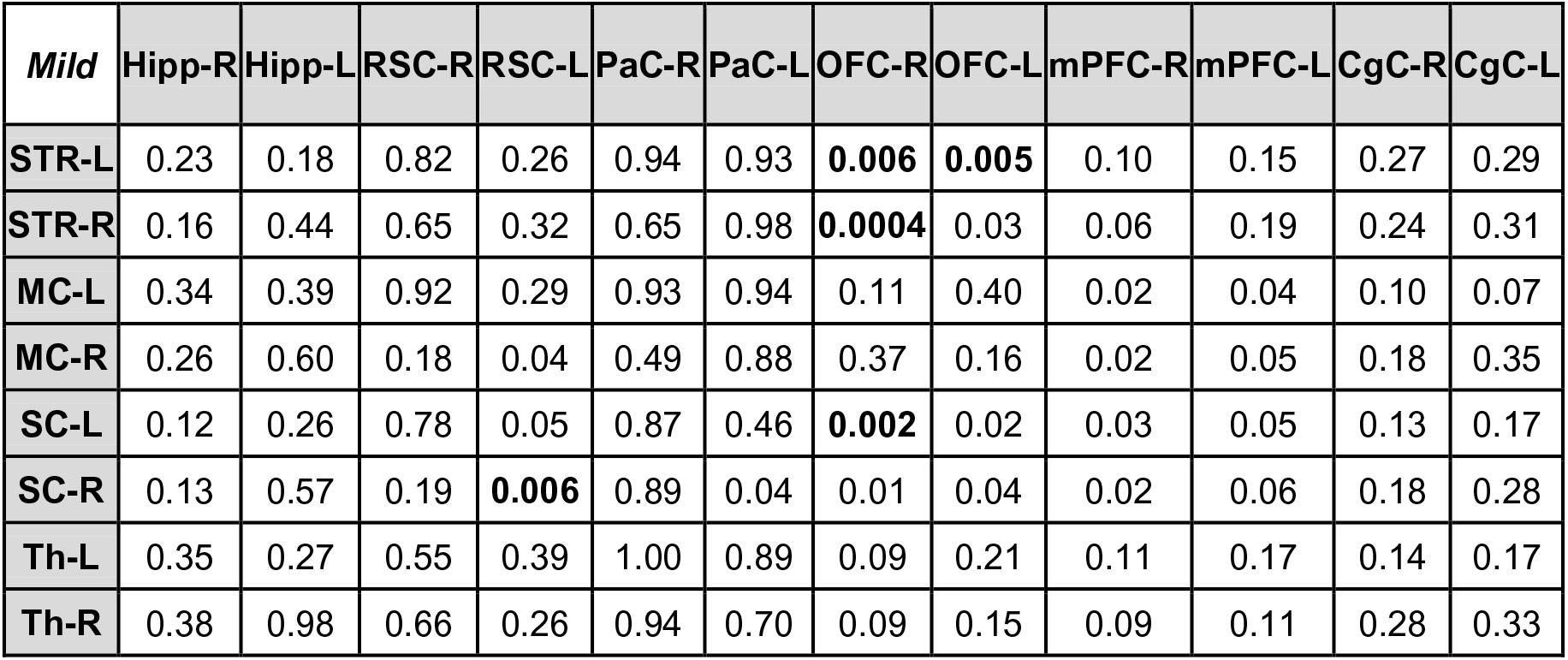
*P* values of the functional connectivity internetwork changes in regions of the DMN and SMN for rats with *mild* VMAT2 knockdown. Data were analyzed using paired t-tests.

**Supplementary Table 6:**
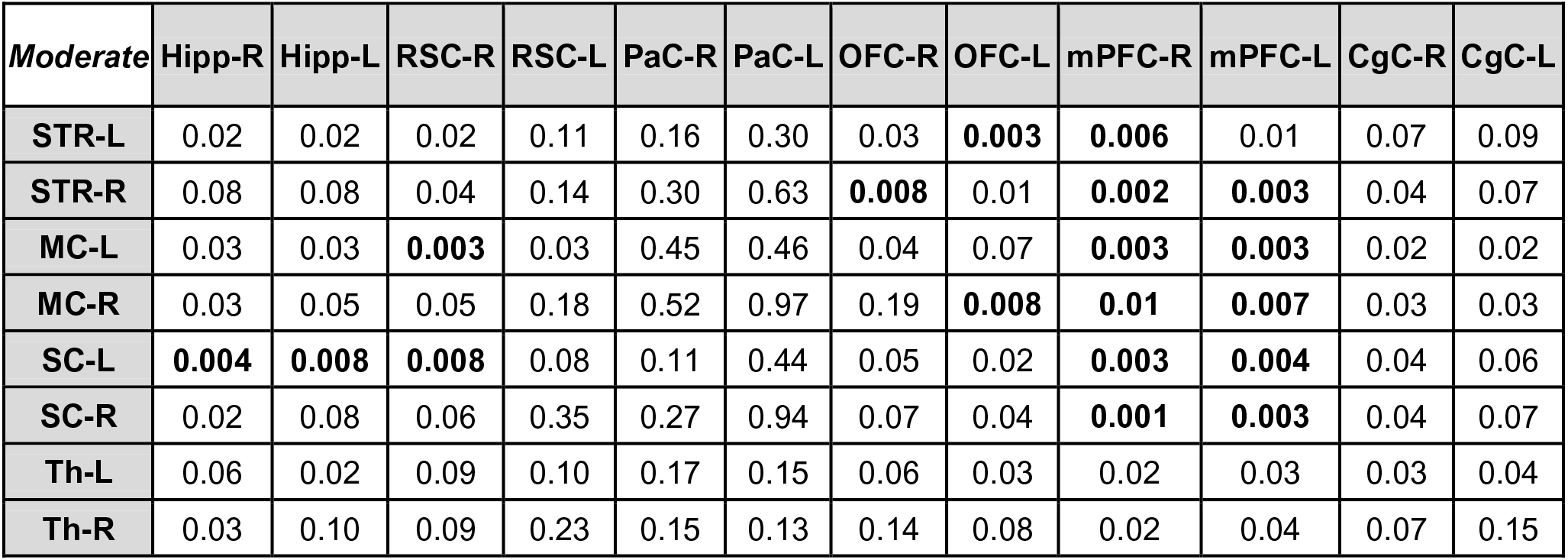
*P* values of the functional connectivity internetwork changes in regions of the DMN and SMN for rats with *moderate* VMAT2 knockdown. Data were analyzed using paired t- tests.

**Supplementary Table 7:**
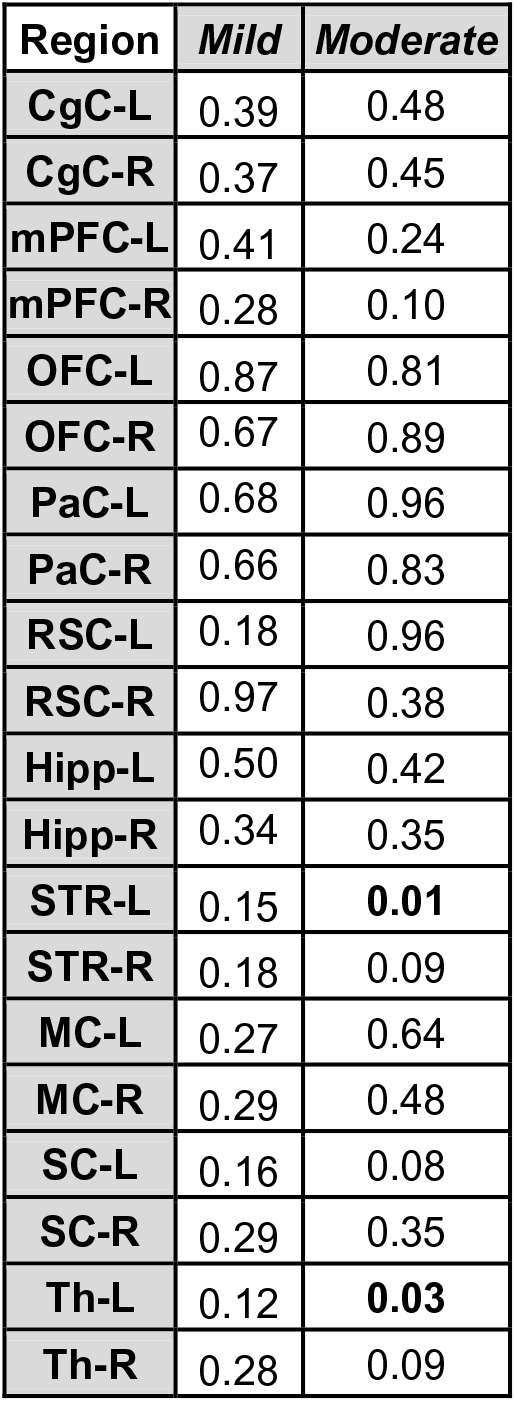
Bonferroni-Sidak corrected *P* values for graph theoretical measures of the global mean connection distance in *mild* and *moderate* VMAT2 knockdown rats. Data were analyzed using paired t-tests.

**Supplementary Table 8:**
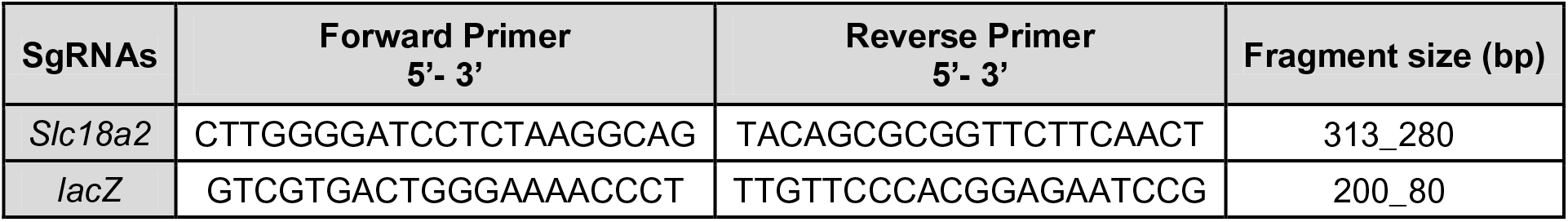
Primers used to amplify the *Slc18a2* and the *lacZ* loci in the Surveyor assay and DNA expected fragments sizes.

**Supplementary Table 9:**
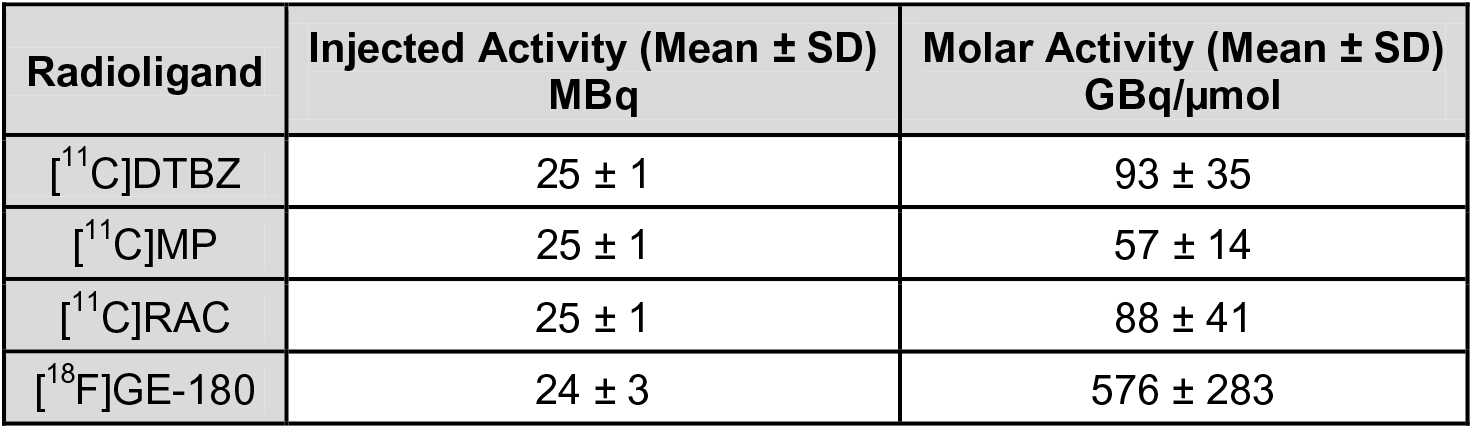
[^11^C]DTBZ, [^11^C]methylphenidate, [^11^C]raclopride and [^18^F]GE-180 injected and molar activities.

**Supplementary Table 10:**
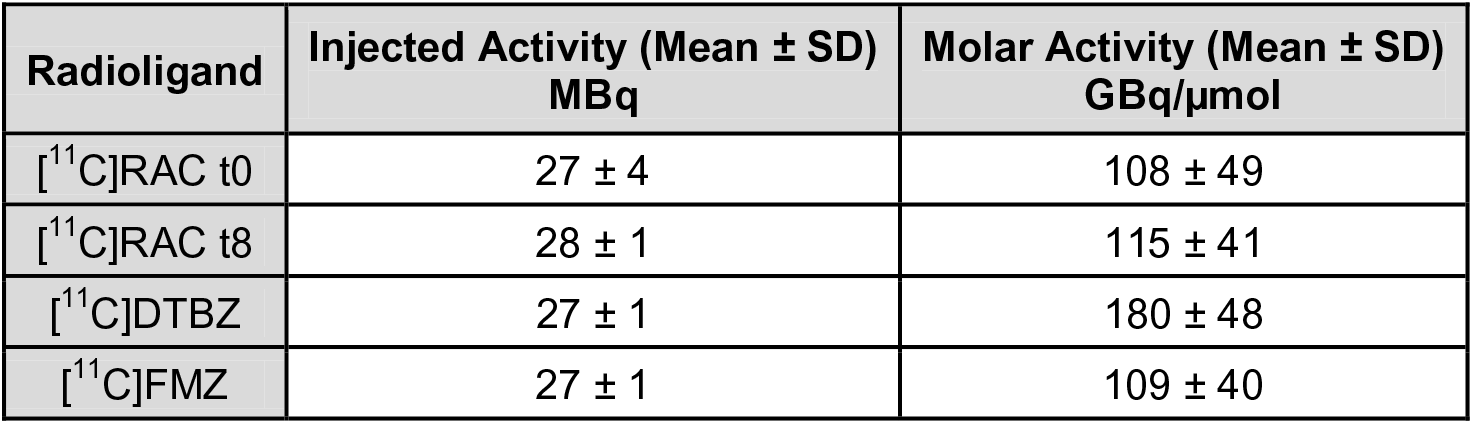
[^11^C]raclopride, [^11^C]DTBZ, [^11^C]flumazenil injected and molar activities.

