## supplemental information for "Combining CRISPR/Cas9 and brain imaging: from genes to molecules to networks"

### Supplementary Figures

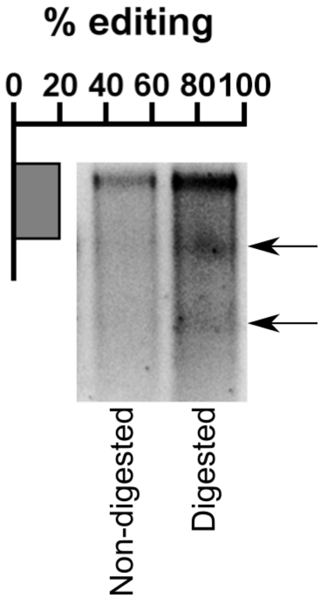


**Supplementary Fig. 1** ***In vitro*** **validation of CRISPR/SaCas9-induced VMAT2 knockdown in rat primary cortical neurons.** One-week post-transduction with conditional vectors for SaCas9 and sgRNA-*Slc18a2*, neurons were processed for the Surveyor assay. The expected cleavage products (arrows), and estimated editing of 20%, could be seen for the digested DNA.

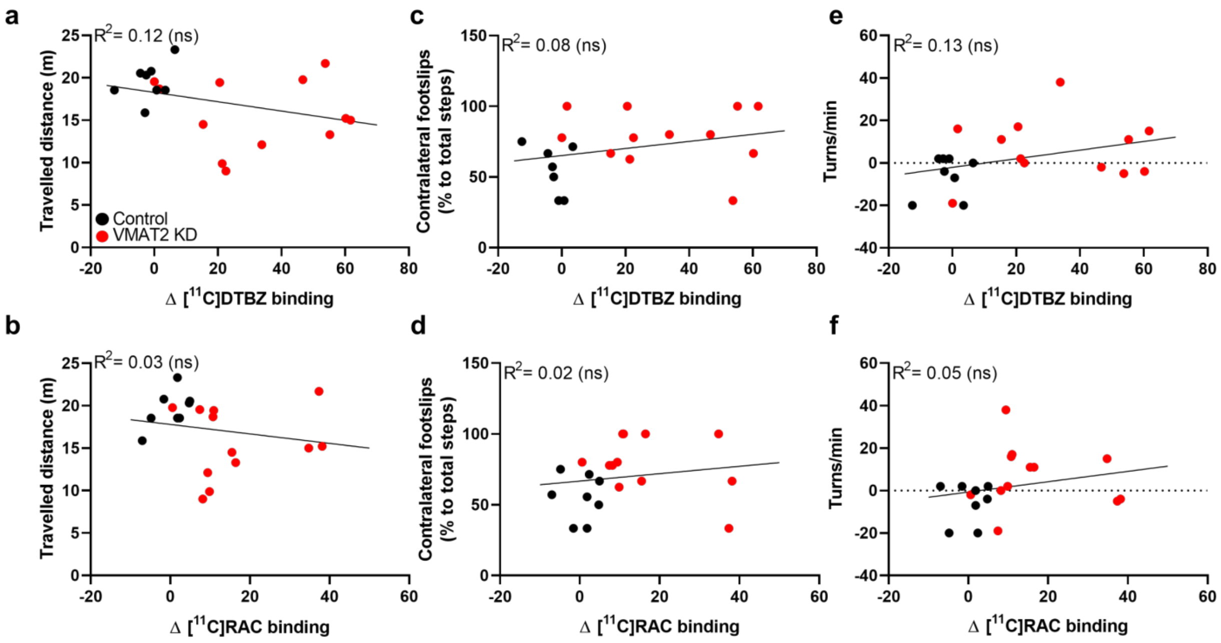


**Supplementary Fig. 2 CRISPR/SaCas9-induced VMAT2 knockdown, postsynaptic changes, and motor impairment do not correlate for whole-body movements.** No correlation was observed between locomotor activity and (**a**) VMAT2 expression changes (Δ [$^{11}$C]DTBZ binding), or (**b**) dopamine availability (Δ [$^{11}$C]RAC binding). The number of contralateral footslips in the beam walk test did not correlate with (**c**) changes in VMAT2 expression (Δ [$^{11}$C]DTBZ binding) or (**d**) dopamine availability (Δ [$^{11}$C]RAC binding). Spontaneous rotations did not correlate with (**e**) changes in VMAT2 expression (Δ [$^{11}$C]DTBZ binding) or (**f**) dopamine availability (Δ [$^{11}$C]RAC binding). ns $P \geq 0.05$. Control rats n= 8; VMAT2 KD rats n= 12. KD, knockdown; [$^{11}$C]RAC, [$^{11}$C]raclopride.

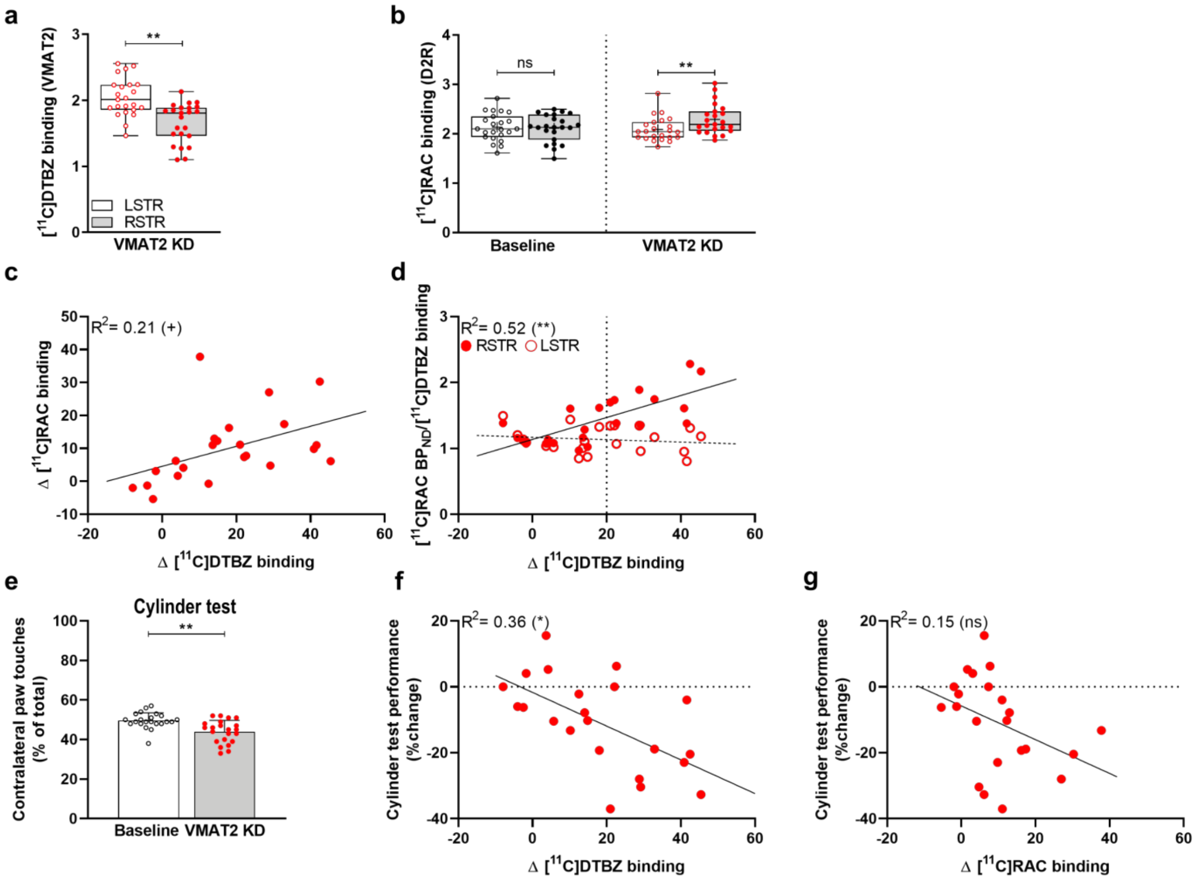


**Supplementary Fig. 3 Reproducibility of the CRISPR/SaCas9-induced VMAT2 knockdown.** (**a**) [$^{11}$C]DTBZ in VMAT2 KD rats and (**b**) [$^{11}$C]RAC PET at baseline and after CRISPR/SaCas9-induced VMAT2 KD. (**c**) A strong correlation between Δ [$^{11}$C]RAC and Δ [$^{11}$C]DTBZ binding is shown. (**d**) At 20% VMAT2 KD (Δ [$^{11}$C]DTBZ binding), D2R binding (Δ [$^{11}$C]RAC binding) prominently increased. This threshold was therefore set to separate the rats into *mild* and *moderate*. Data are shown as boxplot with the median value (central mark), the mean value (plus sign), interquartile range (boxes edges), and the extreme points of the distribution (whiskers). Baseline n= 23, VMAT2 KD n= 23. (**e**) Cylinder test at baseline (n= 22) and 12 - 14 weeks after CRISPR/SaCas9 gene-editing (n= 22). VMAT2 KD rats showed reduced contralateral paw touches compared to baseline. Rats performance in the cylinder test, % change from baseline, correlated with reduced VMAT2 expression (Δ [$^{11}$C]DTBZ binding) (**f**), but not with dopamine availability (Δ [$^{11}$C]RAC binding) (**g**). Data are shown as mean ± SD. $^{+}P< 0.05$, $*P< 0.01$, $**P< 0.001$. KD, knockdown; [$^{11}$C]RAC, [$^{11}$C]raclopride.

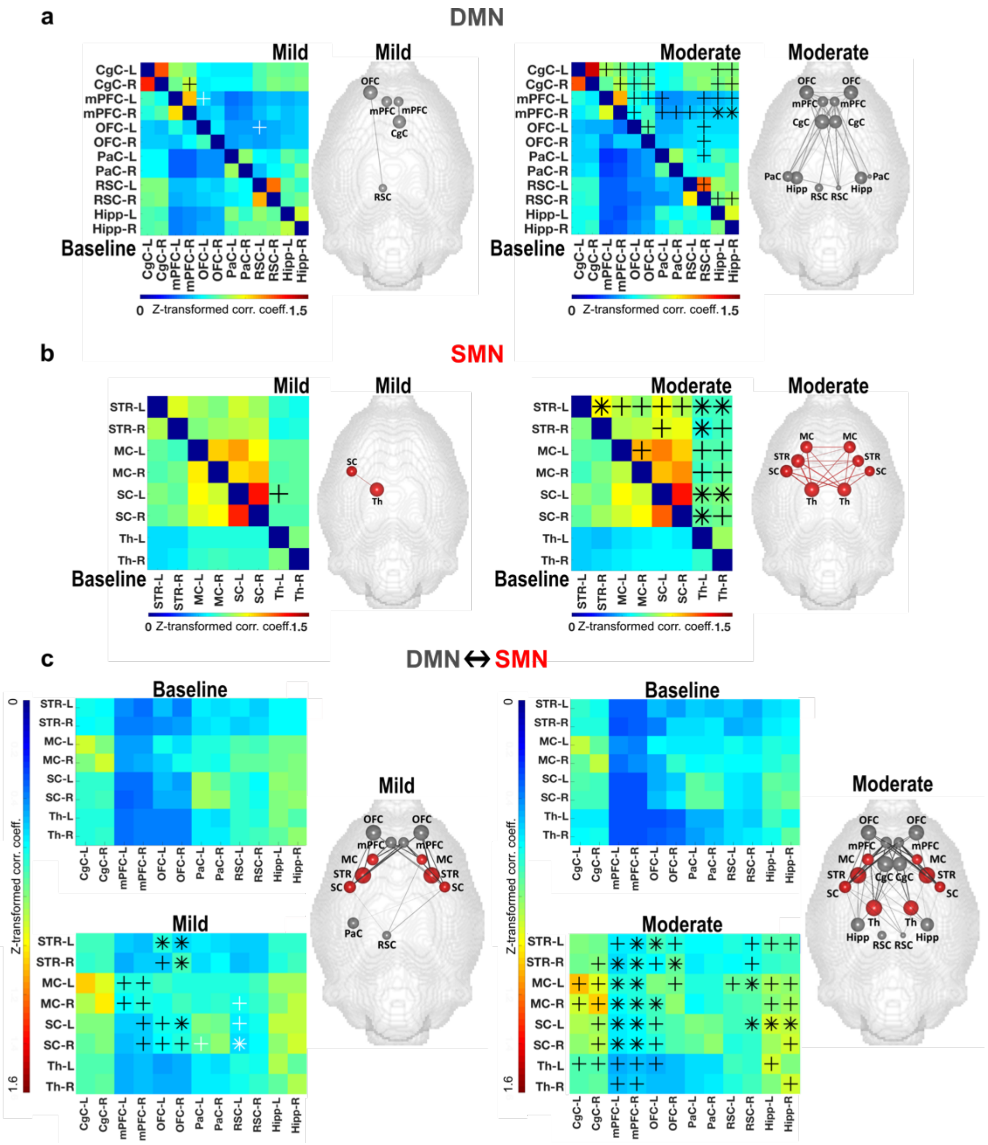


**Supplementary Fig. 4 Increased resting-state functional connectivity after CRISPR/SaCas9-induced VMAT2 knockdown.** Group level correlation matrices of the DMN (**a**) and SMN (**b**) at baseline and after VMAT2 KD for *mild* (left panel) and *moderate* (right panel) rats. Brain graphs, right to the matrices, illustrate the nodes and edges (raw values) that demonstrated rs-FC changes to baseline (%) in DMN (**a**) and SMN (**b**). (**a**) In rats with *mild* VMAT2 KD, we observed divergent rs-FC changes to baseline in regions of the anterior DMN (left mPFC and OFC, right mPFC and CgC). In rats with *moderate* VMAT2 KD, we observed rs-FC increase in most regions of the anterior DMN, extending to the posterior DMN (RSC bilaterally, right RSC, and Hipp bilaterally). (**b**) In rats with *mild* VMAT2 KD, we observed rs-FC increase between the contralateral SC and Th, which intensified in *moderate* VMAT2 KD rats and extended to most of the regions of the SMN. (**c**) Group level correlation matrices of the DMN-SMN rs-FC changes at baseline and after VMAT2 KD for *mild* (left panel) and *moderate* (right panel) rats. Brain graphs, right to the matrices, illustrate the nodes and edges (raw values) that demonstrated internetwork rs-FC changes to baseline (%). Internetwork FC changes in rats with *mild* VMAT2 KD indicated increased rs-FC between anterior regions of the DMN and the SMN (between OFC and STR, SC bilaterally, between right mPFC and MC, SC bilaterally), and decreased rs-FC between posterior regions of the DMN and the SMN (between left RSC and right MC, and SC bilaterally). Increased DMN-SMN rs-FC was found in rats with *moderate* VMAT2 KD. Internetwork rs-FC alterations extended between CgC/Hipp, and SC, MC, Th. $^{+}P < 0.05$, $^{*}P < 0.01$. *Mild*: $\Delta[^{11}C]$DTBZ binding < 20%, n= 13; *Moderate*: $\Delta[^{11}C]$DTBZ binding ≥ 20%, n= 10. KD, knockdown; DMN, default-mode network; SMN, sensorimotor network. mPFC, medial prefrontal cortex; OFC, orbitofrontal cortex; CgC, cingulate cortex; RSC, retrosplenial cortex; Hipp, hippocampus; SC, somatosensory cortex; Th, thalamus; STR, striatum; MC, motor cortex. Abbreviations of brain regions considered for the analysis of the fMRI data, including their respective volumes, are reported in Table 2.

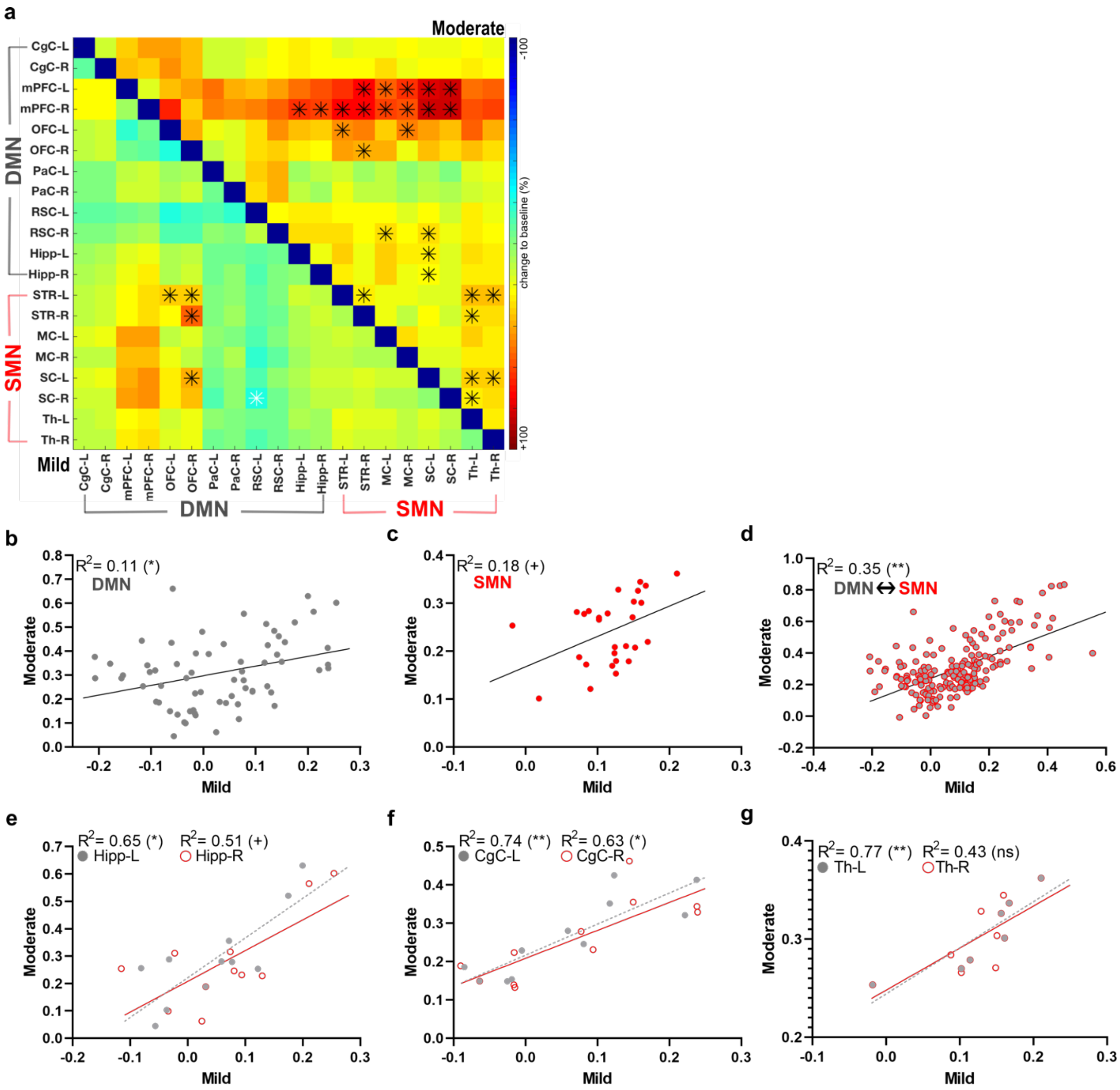


**Supplementary Fig. 5 Node correlation analyses evidence pattern similarity and a linear relationship between the magnitude of the resting-state functional connectivity changes in *mild* and *moderate* rats.** (**a**) Group level correlation matrices of the intraregional and internetwork rs-FC changes to baseline (%) for *mild* and *moderate* VMAT2 KD rats. (**b-d**) Group level intraregional and internetwork rs-FC changes correlated linearly between the two groups. Node correlation analysis between *mild* and *moderate* VMAT2 KD rats indicated a linear increase in the magnitude of the rs-FC changes to baseline in the right and left Hipp (**e**), and CgC (**f**). (**g**) Rs-FC changes (%) in the SMN correlated linearly between *mild* and *moderate* VMAT2 KD rats in the left, but not in the right Th. Our data suggest a similarity in the pattern of the intraregional and internetwork rs-FC changes between rats with *mild* and *moderate* VMAT2 KD and a linear relationship between the magnitude of the rs-FC changes. $^{+}P< 0.05$, $*P< 0.01$, $**P< 0.001$. *Mild*: Δ [$^{11}$C]DTBZ binding < 20%, n= 13; *Moderate*: Δ [$^{11}$C]DTBZ binding ≥ 20%, n= 10. KD, knockdown; DMN, default-mode network; SMN, sensorimotor network. Hipp, hippocampus; CgC, cingulate cortex. Th, thalamus. Abbreviations of brain regions considered for the analysis of the fMRI data, including their respective volumes, are reported in Table 2.

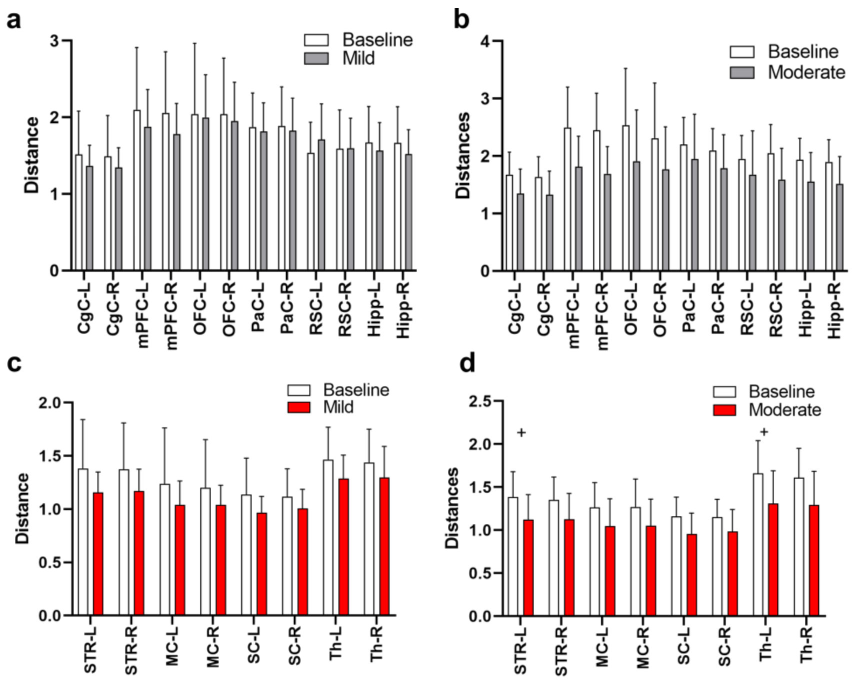


**Supplementary Fig. 6 Graph theoretical analyses reveal lateralized recruitment of SMN regions following the CRISPR/SaCas9-induced VMAT2 *moderate* knockdown.** Quantitative graph theoretical analyses of the global mean connection distance indicated no changes for *mild* (**a**) and *moderate* (**b**) VMAT2 KD rats in the DMN. (**c**) Rats with *mild* VMAT2 KD displayed no changes in the SMN. (**d**) Instead, rats with *moderate* VMAT2 KD presented shorter functional paths to the nodes in the contralateral Th and STR. Data are shown as mean ± SD. $^{+}P < 0.05$, Bonferroni-Sidak-corrected. *Mild*: $\Delta$ [$^{11}$C]DTBZ binding < 20%, n= 13; *Moderate*: $\Delta$ [$^{11}$C]DTBZ binding ≥ 20%, n= 10. KD, knockdown; DMN: default-mode network, SMN: sensorimotor network; Th: Thalamus; STR: striatum. Abbreviations of brain regions considered for the analysis of the fMRI data, including their respective volumes, are reported in Table 2.

### Supplementary Tables

**Supplementary Table 1: *P* values of the functional connectivity changes in regions of the DMN for rats with *mild* VMAT2 knockdown. Data were analyzed using paired t-tests.**

| ***Mild*** | **Hipp-R** | **Hipp-L** | **RSC-R** | **RSC-L** | **PaC-R** | **PaC-L** | **OFC-R** | **OFC-L** | **mPFC-R** | **mPFC-L** | **CgC-R** |
|---|---|---|---|---|---|---|---|---|---|---|---|
| **CgC-L** | 0.40 | 0.56 | 0.95 | 0.33 | 0.76 | 0.84 | 0.22 | 0.27 | 0.07 | 0.07 | 0.61 |
| **CgC-R** | 0.30 | 0.41 | 0.83 | 0.29 | 0.84 | 0.86 | 0.17 | 0.23 | 0.03 | 0.08 | |
| **mPFC-L** | 0.21 | 0.34 | 0.77 | 0.73 | 0.36 | 0.55 | 0.46 | 0.02 | 0.53 | | |
| **mPFC-R** | 0.12 | 0.23 | 0.54 | 0.99 | 0.24 | 0.46 | 0.24 | 0.61 | | | |
| **OFC-L** | 0.49 | 0.55 | 0.20 | 0.05 | 0.51 | 0.67 | 0.18 | | | | |
| **OFC-R** | 0.31 | 0.20 | 0.26 | 0.13 | 0.23 | 0.45 | | | | | |
| **PaC-L** | 0.61 | 0.39 | 0.84 | 0.08 | 0.31 | | | | | | |
| **PaC-R** | 0.74 | 0.59 | 0.83 | 0.11 | | | | | | | |
| **RSC-L** | 0.18 | 0.22 | 0.31 | | | | | | | | |
| **RSC-R** | 0.75 | 0.65 | | | | | | | | | |
| **Hipp-L** | 0.67 | | | | | | | | | | |

**Supplementary Table 2: *P* values of the functional connectivity changes in regions of the DMN for rats with *moderate* VMAT2 knockdown. Data were analyzed using paired t-tests.**

| ***Moderate*** | **Hipp-R** | **Hipp-L** | **RSC-R** | **RSC-L** | **PaC-R** | **PaC-L** | **OFC-R** | **OFC-L** | **mPFC-R** | **mPFC-L** | **CgC-R** |
|---|---|---|---|---|---|---|---|---|---|---|---|
| **CgC-L** | 0.05 | 0.02 | 0.07 | 0.11 | 0.24 | 0.22 | 0.05 | 0.03 | 0.03 | 0.04 | 0.07 |
| **CgC-R** | 0.03 | 0.02 | 0.08 | 0.14 | 0.21 | 0.25 | 0.04 | 0.01 | 0.03 | 0.05 | |
| **mPFC-L** | 0.01 | 0.03 | 0.03 | 0.09 | 0.05 | 0.04 | 0.04 | 0.04 | 0.06 | | |
| **mPFC-R** | **0.005** | **0.007** | 0.01 | 0.03 | 0.03 | 0.03 | 0.06 | 0.01 | | | |
| **OFC-L** | 0.08 | 0.10 | 0.04 | 0.13 | 0.24 | 0.21 | 0.05 | | | | |
| **OFC-R** | 0.16 | 0.11 | 0.03 | 0.07 | 0.29 | 0.47 | | | | | |
| **PaC-L** | 0.35 | 0.60 | 0.03 | 0.10 | 0.13 | | | | | | |
| **PaC-R** | 0.42 | 0.32 | 0.06 | 0.22 | | | | | | | |
| **RSC-L** | 0.10 | 0.07 | 0.02 | | | | | | | | |
| **RSC-R** | 0.03 | 0.04 | | | | | | | | | |
| **Hipp-L** | 0.15 | | | | | | | | | | |

**Supplementary Table 3: *P* values of the functional connectivity changes in regions of the SMN for rats with *mild* VMAT2 knockdown. Data were analyzed using paired t-tests.**

| ***Mild*** | **Th-R** | **Th-L** | **SC-R** | **SC-L** | **MC-R** | **MC-L** | **STR-R** |
|---|---|---|---|---|---|---|---|
| **STR-L** | 0.13 | 0.06 | 0.21 | 0.14 | 0.23 | 0.21 | 0.39 |
| **STR-R** | 0.13 | 0.09 | 0.19 | 0.13 | 0.39 | 0.29 | |
| **MC-L** | 0.24 | 0.16 | 0.47 | 0.16 | 0.46 | | |
| **MC-R** | 0.29 | 0.23 | 0.38 | 0.21 | | | |
| **SC-L** | 0.14 | 0.04 | 0.76 | | | | |
| **SC-R** | 0.30 | 0.12 | | | | | |
| **Th-L** | 0.79 | | | | | | |
| **Th-R** | | | | | | | |

**Supplementary Table 4: *P* values of the functional connectivity changes in regions of the SMN for rats with *moderate* VMAT2 knockdown. Data were analyzed using paired t-tests.**

| ***Moderate*** | **Th-R** | **Th-L** | **SC-R** | **SC-L** | **MC-R** | **MC-L** | **STR-R** |
|---|---|---|---|---|---|---|---|
| **STR-L** | **0.01** | **0.004** | 0.02 | 0.05 | 0.03 | 0.04 | **0.007** |
| **STR-R** | 0.02 | **0.008** | 0.06 | 0.04 | 0.16 | 0.07 | |
| **MC-L** | 0.04 | 0.01 | 0.10 | 0.08 | 0.04 | | |
| **MC-R** | 0.02 | 0.02 | 0.07 | | | | |
| **SC-L** | **0.003** | **0.001** | 0.11 | | | | |
| **SC-R** | 0.02 | **0.008** | | | | | |
| **Th-L** | 0.07 | | | | | | |
| **Th-R** | | | | | | | |

**Supplementary Table 5: *P* values of the functional connectivity internetwork changes in regions of the DMN and SMN for rats with *mild* VMAT2 knockdown. Data were analyzed using paired t-tests.**

| ***Mild*** | **Hipp-R** | **Hipp-L** | **RSC-R** | **RSC-L** | **PaC-R** | **PaC-L** | **OFC-R** | **OFC-L** | **mPFC-R** | **mPFC-L** | **CgC-R** | **CgC-L** |
|---|---|---|---|---|---|---|---|---|---|---|---|---|
| **STR-L** | 0.23 | 0.18 | 0.82 | 0.26 | 0.94 | 0.93 | **0.006** | **0.005** | 0.10 | 0.15 | 0.27 | 0.29 |
| **STR-R** | 0.16 | 0.44 | 0.65 | 0.32 | 0.65 | 0.98 | **0.0004** | 0.03 | 0.06 | 0.19 | 0.24 | 0.31 |
| **MC-L** | 0.34 | 0.39 | 0.92 | 0.29 | 0.93 | 0.94 | 0.11 | 0.40 | 0.02 | 0.04 | 0.10 | 0.07 |
| **MC-R** | 0.26 | 0.60 | 0.18 | 0.04 | 0.49 | 0.88 | 0.37 | 0.16 | 0.02 | 0.05 | 0.18 | 0.35 |
| **SC-L** | 0.12 | 0.26 | 0.78 | 0.05 | 0.87 | 0.46 | **0.002** | 0.02 | 0.03 | 0.05 | 0.13 | 0.17 |
| **SC-R** | 0.13 | 0.57 | 0.19 | **0.006** | 0.89 | 0.04 | 0.01 | 0.04 | 0.02 | 0.06 | 0.18 | 0.28 |
| **Th-L** | 0.35 | 0.27 | 0.55 | 0.39 | 1.00 | 0.89 | 0.09 | 0.21 | 0.11 | 0.17 | 0.14 | 0.17 |
| **Th-R** | 0.38 | 0.98 | 0.66 | 0.26 | 0.94 | 0.70 | 0.09 | 0.15 | 0.09 | 0.11 | 0.28 | 0.33 |

**Supplementary Table 6: *P* values of the functional connectivity internetwork changes in regions of the DMN and SMN for rats with *moderate* VMAT2 knockdown. Data were analyzed using paired t-tests.**

| ***Moderate*** | **Hipp-R** | **Hipp-L** | **RSC-R** | **RSC-L** | **PaC-R** | **PaC-L** | **OFC-R** | **OFC-L** | **mPFC-R** | **mPFC-L** | **CgC-R** | **CgC-L** |
|---|---|---|---|---|---|---|---|---|---|---|---|---|
| **STR-L** | 0.02 | 0.02 | 0.02 | 0.11 | 0.16 | 0.30 | 0.03 | **0.003** | **0.006** | 0.01 | 0.07 | 0.09 |
| **STR-R** | 0.08 | 0.08 | 0.04 | 0.14 | 0.30 | 0.63 | **0.008** | 0.01 | **0.002** | **0.003** | 0.04 | 0.07 |
| **MC-L** | 0.03 | 0.03 | **0.003** | 0.03 | 0.45 | 0.46 | 0.04 | 0.07 | **0.003** | **0.003** | 0.02 | 0.02 |
| **MC-R** | 0.03 | 0.05 | 0.05 | 0.18 | 0.52 | 0.97 | 0.19 | **0.008** | **0.01** | **0.007** | 0.03 | 0.03 |
| **SC-L** | **0.004** | **0.008** | **0.008** | 0.08 | 0.11 | 0.44 | 0.05 | 0.02 | **0.003** | **0.004** | 0.04 | 0.06 |
| **SC-R** | 0.02 | 0.08 | 0.06 | 0.35 | 0.27 | 0.94 | 0.07 | 0.04 | **0.001** | **0.003** | 0.04 | 0.07 |
| **Th-L** | 0.06 | 0.02 | 0.09 | 0.10 | 0.17 | 0.15 | 0.06 | 0.03 | 0.02 | 0.03 | 0.03 | 0.04 |
| **Th-R** | 0.03 | 0.10 | 0.09 | 0.23 | 0.15 | 0.13 | 0.14 | 0.08 | 0.02 | 0.04 | 0.07 | 0.15 |

**Supplementary Table 7: Bonferroni-Sidak corrected *P* values for graph theoretical measures of the global mean connection distance in *mild* and *moderate* VMAT2 knockdown rats. Data were analyzed using paired t-tests.**

| **Region** | ***Mild*** | ***Moderate*** |
|---|---|---|
| **CgC-L** | 0.39 | 0.48 |
| **CgC-R** | 0.37 | 0.45 |
| **mPFC-L** | 0.41 | 0.24 |
| **mPFC-R** | 0.28 | 0.10 |
| **OFC-L** | 0.87 | 0.81 |
| **OFC-R** | 0.67 | 0.89 |
| **PaC-L** | 0.68 | 0.96 |
| **PaC-R** | 0.66 | 0.83 |
| **RSC-L** | 0.18 | 0.96 |
| **RSC-R** | 0.97 | 0.38 |
| **Hipp-L** | 0.50 | 0.42 |
| **Hipp-R** | 0.34 | 0.35 |
| **STR-L** | 0.15 | **0.01** |
| **STR-R** | 0.18 | 0.09 |
| **MC-L** | 0.27 | 0.64 |
| **MC-R** | 0.29 | 0.48 |
| **SC-L** | 0.16 | 0.08 |
| **SC-R** | 0.29 | 0.35 |
| **Th-L** | 0.12 | **0.03** |
| **Th-R** | 0.28 | 0.09 |

**Supplementary Table 8: Primers used to amplify the *Slc18a2* and the *lacZ* loci in the Surveyor assay and DNA expected fragments sizes.**

| SgRNAs | Forward Primer 5'- 3' | Reverse Primer 5'- 3' | Fragment size (bp) |
|---|---|---|---|
| *Slc18a2* | CTTGGGGATCCTCTAAGGCAG | TACAGCGCGGTTCTTCAACT | 313_280 |
| *lacZ* | GTCGTGACTGGGAAAACCCT | TTGTTCCCACGGAGAATCCG | 200_80 |

**Supplementary Table 9: [$^{11}$C]DTBZ, [$^{11}$C]methylphenidate, [$^{11}$C]raclopride and [$^{18}$F]GE-180 injected and molar activities.**

| Radioligand | Injected Activity (Mean ± SD) MBq | Molar Activity (Mean ± SD) GBq/µmol |
|---|---|---|
| [$^{11}$C]DTBZ | 25 ± 1 | 93 ± 35 |
| [$^{11}$C]MP | 25 ± 1 | 57 ± 14 |
| [$^{11}$C]RAC | 25 ± 1 | 88 ± 41 |
| [$^{18}$F]GE-180 | 24 ± 3 | 576 ± 283 |

**Supplementary Table 10: [$^{11}$C]raclopride, [$^{11}$C]DTBZ, [$^{11}$C]flumazenil injected and molar activities.**

| Radioligand | Injected Activity (Mean ± SD) MBq | Molar Activity (Mean ± SD) GBq/µmol |
|---|---|---|
| [$^{11}$C]RAC t0 | 27 ± 4 | 108 ± 49 |
| [$^{11}$C]RAC t8 | 28 ± 1 | 115 ± 41 |
| [$^{11}$C]DTBZ | 27 ± 1 | 180 ± 48 |
| [$^{11}$C]FMZ | 27 ± 1 | 109 ± 40 |
